# Molecular Basis and Cellular Effects of Janus-Class–Driven Cytoplasmic PYK2 Coacervates

**DOI:** 10.1101/2025.06.24.661287

**Authors:** Giovanni Colombo, Israa Salem, Kacper Szczepski, Piao Yu, Shaden Alfaiyz, Francisco Javier Guzmán-Vega, Ahmed Abogosh, Maxat Kulmanov, Samah AlHarthi, Gress Kadaré, Robert Hoehndorf, Jean-Antoine Girault, Łukasz Jaremko, Afaque A. Momin, Stefan T. Arold

**Affiliations:** KAUST Center of Excellence for Smart Health, Biological and Environmental Science and Engineering Division, King Abdullah University of Science and Technology (KAUST), Thuwal 23955-6900, Kingdom of Saudi Arabia; Genetics section, Research Department, Natural and Health Sciences Research Center, Princess Nourah bint Abdulrahman University, P.O. Box 84428, Riyadh 11671, Saudi Arabia; KAUST Center of Excellence for Smart Health, Computer, Electrical, and Mathematical Science and Engineering Division, King Abdullah University of Science and Technology (KAUST), Thuwal 23955-6900, Kingdom of Saudi Arabia; Inserm UMR-S 1270, Sorbonne Université, Faculty of Sciences and Engineering, Institut du Fer à Moulin, 75005, Paris, France

**Keywords:** focal adhesion, phase separation, cancer, kinase, ESM2, Janus class

## Abstract

Kinase activity is increasingly associated with biomolecular phase separation. Focal adhesion kinase (FAK) forms membrane-associated condensates with paxillin to promote adhesion. Here we show that its paralogue, proline-rich tyrosine kinase 2 (PYK2), undergoes phase separation *via* a distinct mechanism. PYK2 forms cytoplasmic condensates primarily driven by its kinase– FAT linker (KFL) region. Overexpression of PYK2 induces condensates enriched in its autophosphorylated form, which sequester paxillin from focal adhesions and impair cell adhesion. We uncover an autoregulatory mechanism involving the KFL, linking self-association, autophosphorylation, and condensation. Uncommon among known phase separation drivers, KFL condensation is phosphorylation-independent and its sequence belongs to the “Janus” class. Using a transformer-based protein language model, we identified non-homologous sequences with similar features, many from adhesion and cytoskeletal regulators. We validated the phase-separating potential of several of these sequences in cells. These findings reveal a novel mechanism linking phase separation with kinase activation, and demonstrate distinct condensation behavior in close homologues. Our results also highlight how protein concentration modulates condensate function, with implications for disease, and expand the landscape of phase separation drivers.

## INTRODUCTION

The precise spatial and temporal control of kinases is central to faithful signal transduction in normal cells, and kinase dysregulation is associated with cancer and other diseases^[1]^. Liquid-liquid phase separation (LLPS) has emerged as a key mechanism for regulating signalling events by selectively enriching relevant molecules within membrane-less signalosomes, while excluding others^[2]^. By bringing reacting components together, biomolecular condensates enhance the specificity of enzymes, substrates, cofactors, and adaptor proteins, which, otherwise, may be insufficiently selective to promote non-erroneous signalling^[2,3,4,5,6]^. These features make phase separation a suitable mechanism for the spatiotemporal control of kinases. Indeed, kinases have been shown to be involved in several biomolecular condensates, including the aggrephagy machinery, centriole biogenesis, integrin-mediated adhesion, T-cell receptor signalling, and signalling through receptor tyrosine kinases^[2,7,8,9,10,11]^. Despite the growing body of evidence linking kinase functions with LLPS, still very little is known about the molecular mechanisms that control the participation of kinases in condensates, and the effect of condensates on their catalytic activity.

The focal adhesion kinase (FAK) and its close paralogue the proline-rich tyrosine kinase 2 (PYK2) control cellular adhesion, migration, and signalling by orchestrating the assembly and disassembly of integrin adhesion complexes (IACs)^[12,13]^. For FAK, the better studied paralogue, it has been shown that its capacity to phase separate synergistically with other components of focal adhesions (FAs, including p130Cas and paxillin) is essential for initiating the formation of IACs at sites where integrins cluster^[9,14]^. Additional literature has further established the role of biomolecular coacervates in IACs and other adhesion structures^[9,14,15,16]^.

PYK2 and FAK share the same domain architecture consisting of a 4.1, ezrin, radixin, moesin (FERM) domain, a tyrosine kinase domain, and a focal adhesion targeting (FAT) domain (residues 34-365; 418-699; 870-1009, respectively, in PYK2 numbering; **Fig. 1A**). All three domains and the two extended linker regions between them display numerous protein-protein interaction sites. Several of these interaction sites are conserved between PYK2 and FAK, such as an SH2-binding phosphotyrosine (Y402 and Y397 in PYK2 and FAK, respectively), and three SH3-interacting proline-rich motifs (PR1: 377–380; PR2: 713–726; PR3: 855-861; PYK2 numbering. **Fig. 1A**)^[3,17,18,19]^. PYK2 and FAK have overlapping cellular functions, for example, the control of integrin-mediated migration and cellular survival following detachment^[20,21]^. Their capability of promoting cell migration and survival makes PYK2 and FAK key players in the development and wound healing^[22]^. In line with these functions, PYK2 and FAK dysregulation is also associated with both cancer and neurological disorders^[23,24,25]^. In particular, PYK2 plays a critical role in several aspects of cancer progression, including cell migration and invasion^[26,27]^. The degree of PYK2 overexpression can vary depending on the specific type of cancer and many other factors^[26]^. Additionally, PYK2 may play a role in the tumour microenvironment and metastasis^[27]^. These lines of evidence identify PYK2 as an attractive target for antitumor therapies, with a significant functional redundancy to FAK.

**Fig. 1:**
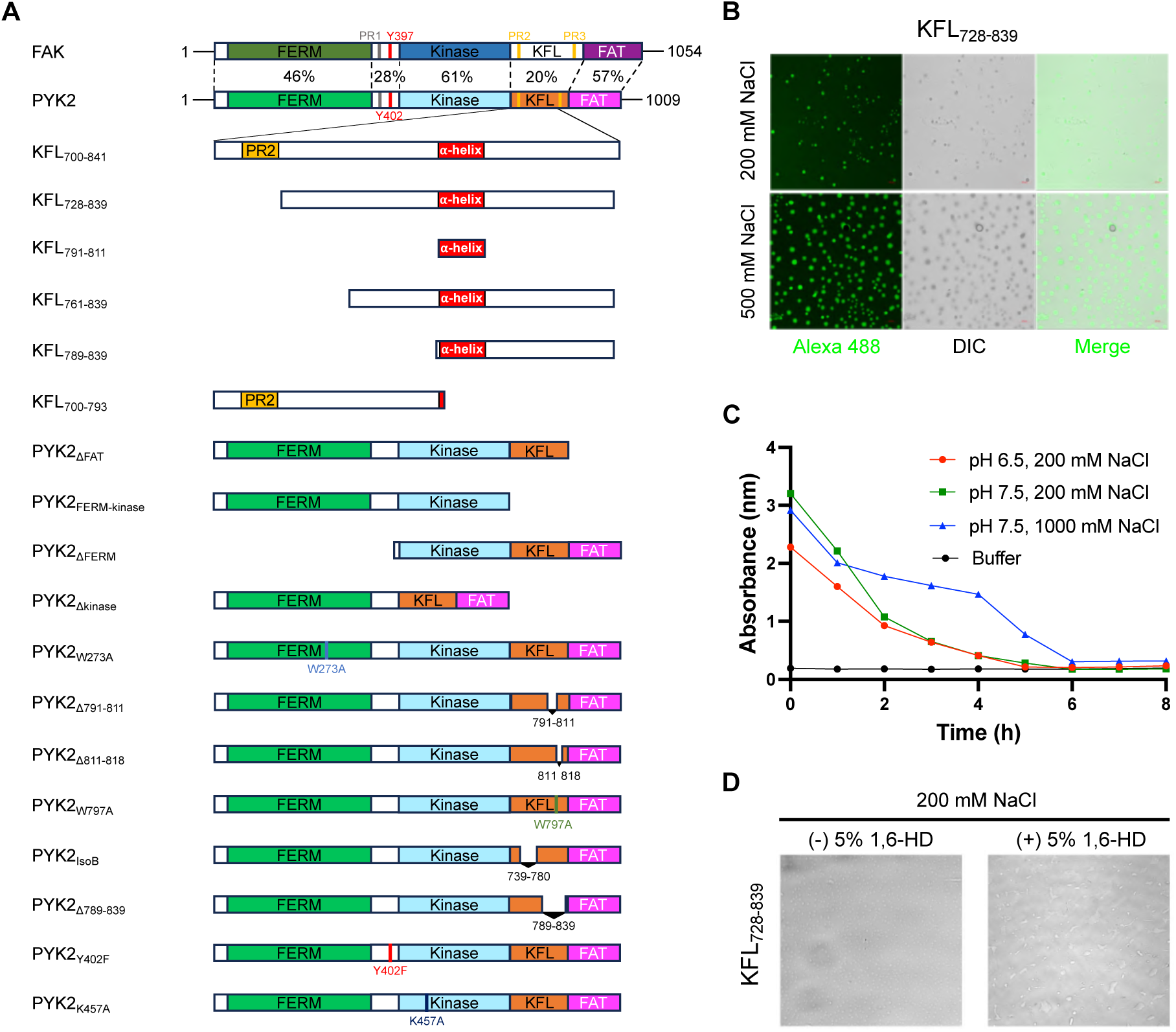
The PYK2 KFL forms droplets *in vitro*. **A.** Schematic representation of FAK and PYK2 constructs with annotated residue numbers. **B.** Widefield fluorescence and DIC images of Alexa 488-labeled KFL_728-839_ with 200 mM (*top row*) and 500 mM NaCl (*bottom row*), showing droplets formation. **C.** OD600 turbidity measurement over time at pH 6.5, 200 mM NaCl (red line); pH 7.5, 200 mM NaCl (green line); pH 7.5 1000 mM NaCl (blue line). **D.** DIC images of non-labeled KFL_728-839_ with 200 mM NaCl in the absence and presence of 5% 1,6-HD. DIC: differential interference contrast. 1,6-HD: 1,6-hexanediol.

Many of the mechanistic aspects and differences in the cellular functions of PYK2 and FAK remain insufficiently understood. Differently from FAK, PYK2 has acquired the capacity to bind calmodulin (CaM), endowing it with the ability to sense and react to calcium influx^[13,28,29,30]^. There is also evidence for PYK2 being less strongly recruited to IACs than FAK, despite containing a paxillin-binding FAT domain that has been shown to mediate recruitment of FAK to IACs^[31,32,33]^. Indeed, PYK2 mainly displays a cytoplasmic localization, possibly in part due to the inability of PYK2’s FAT domain to interact with talin, a key adaptor protein anchoring FAK to FAs^[34]^. Understanding the idiosyncratic differences between PYK2 and FAK is important for discerning their cellular roles, and for efficiently targeting them by therapeutic approaches.

Here we show that PYK2 promotes phase separation *in vitro* and in cells. However, despite its overall similarity to FAK, PYK2 phase separates in the cytoplasm, rather than at sites of cellular adhesions. Combining biochemical, structural, and cellular analyses, we identify the molecular mechanisms underlying PYK2 phase separation, and explore the cellular effects of these condensates. Using AI-based methods, we identify additional driver regions of the same class in other proteins involved in adhesion and cytoskeletal organisation, extending the functional landscape of sequences associated with phase separation.

## RESULTS

### The PYK2 KFL forms condensates in vitro

Previously, we showed that the PYK2 kinase-FAT linker (KFL) region between PR2 and PR3 (KFL_728-839_), which is disordered except for an α-helix (residues 791-811), forms dimers *in vitro* that interact with CaM^[30]^. KFL_728-839_ remains highly flexible in these interactions^[30]^. During our work with recombinant KFL_728-839_, we noticed the appearance of liquid-like droplets *in vitro*. The formation of homotypic condensates by KFL_728-839_ was unexpected because as an apparent dimer, it lacked the multivalency that is generally considered a requirement for LLPS^[35,36]^. However, we consistently observed droplets in images acquired using differential interference contrast (DIC) and widefield fluorescence microscopy of fluorescently labelled recombinant and purified KFL_728-839_ (**Fig. 1B**). Droplet formation increased with increases in the concentration of protein and NaCl (**Fig. 1B** and **Suppl. Table 1**), but decreased when the pH dropped from 7.5 to 6.5 (**Fig. 1C** and **Suppl. Fig. 1A**). The addition of 5% 1,6-hexanediol (1,6-HD), an alcohol that disrupts weak hydrophobic interactions of proteins, substantially impaired droplet formation, providing a hallmark feature of LLPS (**Fig. 1D**). Shorter fragments of PYK2 KFL (residues 761–839 and 789–839) failed to form droplets under the same conditions (**Suppl. Fig. 1B**). These shorter PYK2 KFL fragments also do not dimerise^[30]^, suggesting that dimerisation is required for LLPS.

We also previously observed that the KFL from FAK is able to form homodimers, but not heterodimers with the KFL of PYK2^[30]^. Therefore, we tested FAK KFL fragments for droplet formation *in vitro*. Neither FAK KFL_776–841_ (which dimerises) nor KFL_804–832_ (which does not dimerise)^[30,37]^ formed visible droplets (**Suppl. Fig. 1B**). A longer FAK KFL fragment (residues 764-845) aggregated under the *in vitro* conditions tested^[30]^. Thus, under the conditions used, the KFL of PYK2 but not of FAK was sufficient to form droplets *in vitro*.

### The LLPS transition only entails subtle chemical environment changes in KFL

To probe the molecular basis of KFL_728-839_ phase separation, we used NMR spectroscopy to compare the chemical environments of residues in the dilute (NoLLPS) and dense (LLPS) phases. We compared an LLPS-containing KFL_728-839_ sample, measured after 24 hours incubation in the NMR tube, to the same sample titrated with LLPS-disrupting 5% 1,6-HD. This procedure only introduced several minor residue-specific chemical shift perturbations (CSPs) (**Fig. 2A, B**). Differences between LLPS/NoLLPS samples observed in the regions of Cα/CO and Cβ/CO in the ^13^C detected CBCACO spectra were significant (i.e. above the experimental/digital resolution of the identified signal positions), but too small to indicate conformational or angular changes in side chains. The signal shape and intensity of the CBCACO correlations remained consistent between LLPS and NoLLPS conditions, with no evidence of line broadening, suggesting that side-chain dynamics were largely unaffected. Thus, the transition between LLPS and NoLLPS conditions after addition of 1,6-HD only led to subtle chemical environment changes around the CSP displaying atoms/residues. Residues displaying these subtle changes were enriched in the helical region of KFL_728-839_ (**Fig. 2A, B, C** and **Suppl. Fig. 1C**), previously identified as a key site for homodimerisation and CaM binding^[30]^. These findings support a model in which homotypic KFL_728-839_ condensation results from dimer-like fuzzy intermolecular interactions between one molecule and additional molecules in its vicinity. Thus, conceptually, condensates appear to be a fuzzy multivalent extension of the fuzzy KFL_728-839_ dimers.

**Fig. 2:**
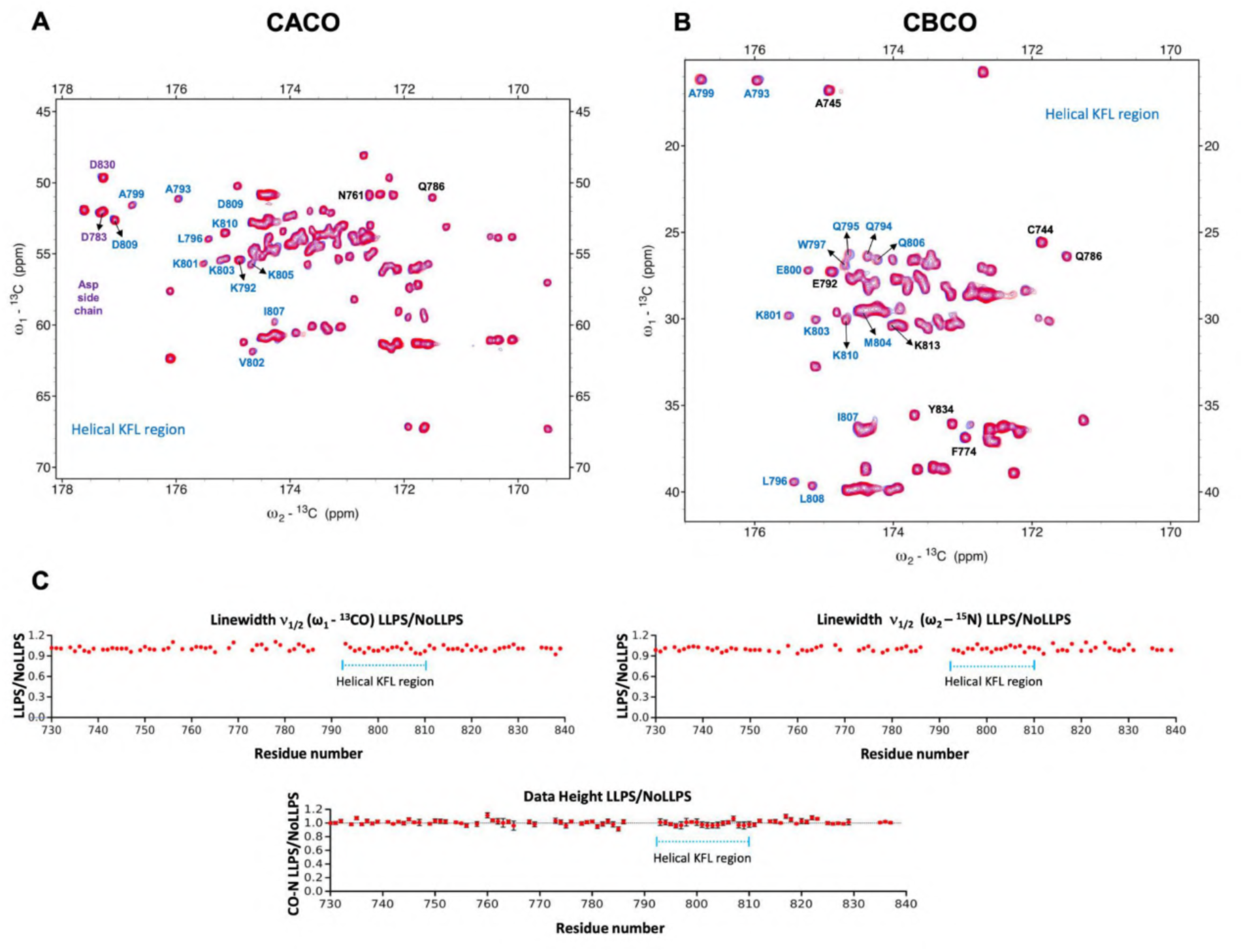
NMR reveals involvement of helical region side chains in PYK2 condensation. Differences between LLPS/NoLLPS samples observed in the regions of Ca/CO (**A.**) and Cb/CO (**B.**) in the ^13^C detected CBCACO spectra reveal distinct conformational alterations in side chains of the helical region of KFL (residues 792-810) while dynamics (**C.**) of the backbone is virtually not affected. Residues exhibiting significant chemical shift changes are highlighted in the spectrum with residues belonging to the helical region colored in blue.

### In vitro condensation of the KFL is not altered by phosphorylation

Phosphorylation of intrinsically disordered proteins can affect their propensity to undergo phase separation^[38]^. In the case of PYK2, its phosphorylation on Y402 recruits and activates Src kinases, leading to phosphorylation of PYK2 KFL by Src kinases and, subsequently, by downstream MAP kinases^[30,39]^. To test whether phosphorylation of the PYK2 KFL alters its LLPS propensity, we incubated purified KFL with purified active kinases (either Src or a constitutively active homologue of the Erk MAP kinase, MPK4). Subsequent mass spectrometry analysis confirmed the *in vitro* phosphorylation of KFL at canonical sites for Src (Y819QWL) and MAP kinases (S752P, S759P, T765P) (**Suppl. Fig. 2A**). However, phosphorylated KFL did not show a significantly altered propensity to form condensates *in vitro* (**Fig. 3A**). Indeed, the phosphorylated residues were neither within the KFL helix, nor within the previously mapped regions involved in dimerisation, nor among those with significant CSPs upon transitioning to LLPS (**Fig. 2A**).

**Fig. 3:**
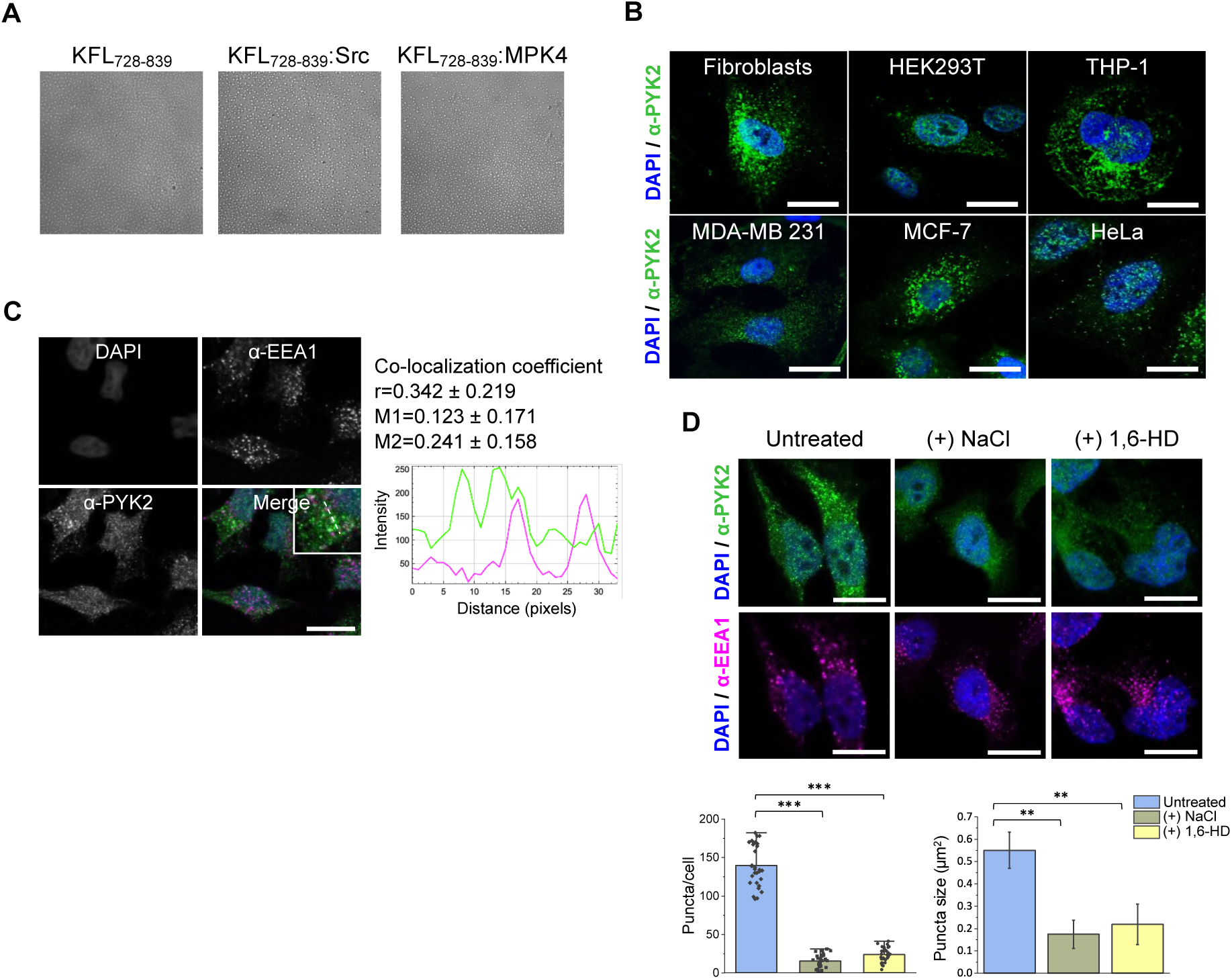
PYK2 forms intracellular puncta responding to biophysical perturbations. **A.** Non-labeled KFL_728-839_, KFL_728-839_:Src and KFL_728-839_:MPK4 droplets visualized by DIC. **B.** Immunofluorescence of endogenous PYK2 (green) in human fibroblasts, HEK293T, THP-1, MDA-MB-231, MCF-7 and HeLa cell lines. Nuclei are counterstained with DAPI (blue). Scale bar: 10 µm. **C.** Single-channel and merged immunofluorescence images of HeLa cells stained with DAPI (blue), anti-EEA1 (magenta), and anti-PYK2 (green) antibodies. Co-localization analysis includes Pearson/Manders (r/M1,M2) coefficients calculated from *n* = 5 and profile plot. Scale bar: 10 µm. **D.** Immunofluorescence images of HeLa cells untreated or treated with NaCl or 1,6-HD (same field of view). Cells were stained with anti-PYK2 (green, *top row*) and anti-EEA1 (magenta, *bottom row*) antibodies and counterstained with DAPI. Bar plots (*left*) show the number of PYK2 puncta per cell; bar plots (*right*) show PYK2 puncta size under each condition. **p < 0.01, ***p < 0.001 compared to untreated. EEA1: early endosomal antigen 1.

### Endogenous PYK2 forms cytoplasmic puncta that are distinct from endosomes

We next investigated the presence and localisation of PYK2 condensates in cells. Confocal microscopy analysis using a PYK2 monoclonal antibody showed that endogenous PYK2 formed distinct cytoplasmic puncta in both normal and cancerous human cell lines, including fibroblasts, HEK293T, THP-1, MCF-7, MDA-MB-231, and HeLa (**Fig. 3B**). Despite notable differences in PYK2 expression levels among these cell types, puncta were consistently observed. Given that PYK2 was originally described as a cytoplasmatic kinase^[31]^ and that is capable of nuclear shuttling^[33]^, we consider it reasonable to conclude that these puncta localize predominantly to perinuclear regions, but not to FAs (**Fig. 3B**). These morphological and distribution patterns were reminiscent of proteins forming dense phases^[40]^.

Previously, PYK2 has been associated with intra-vesicular trafficking, in particular overlapping with early endosomes in certain contexts^[41]^. To rule out the possibility that those puncta were early endosomal structures, PYK2 was immunostained alongside the marker EEA1, and quantitative colocalization was assessed using Pearson’s and Manders’ correlation coefficients (**Fig. 3C**). The Pearson correlation coefficient was 0.342 ± 0.219, indicating a moderate correlation between PYK2 and EEA1 stainings. However, Manders’ coefficients (M1 = 0.123 ± 0.171, M2 = 0.241 ± 0.158) revealed that only a small fraction (∼20%) of PYK2 overlapped with EEA1, suggesting limited direct colocalization (**Fig. 3C**). Moreover, increasing salt concentration in cell culture medium or treatment with 1,6-HD effectively disrupted endogenous PYK2 puncta without affecting EEA1 staining (**Fig. 3D**), suggesting that PYK2 assemblies are maintained through weak intermolecular interactions, likely *via* LLPS. These results indicate that while PYK2 and EEA1 might be present in proximal subcellular regions, PYK2 puncta are structurally distinct from early endosomal vesicles.

### PYK2 expression levels affect PYK2 condensate features

To explore the dynamic behaviour of PYK2 within the cell, we ectopically expressed EGFP-tagged PYK2 in HEK293T cells. EGFP-PYK2 formed cytoplasmic puncta after 24 hours (**Fig. 4A** and **Suppl. Fig. 2B, C**). The size and subcellular location of PYK2 condensates correlated with the amount of plasmids transfected, and hence with cytoplasmic protein concentration. Transfection with 250 ng/cm^2^ of EGFP-PYK2 plasmid resulted in small and dispersed puncta, whereas transfection with ∼900 ng/cm^2^ of plasmid produced puncta with diameters up to 2 µm that were larger than endogenous puncta (**Fig. 4A** and **Suppl. Figure 2D**). Further increasing the plasmid amount led to the occurrence of irregular aggregates (**Suppl. Fig. 2D)**. As observed for endogenous PYK2, droplets formed either by EGFP-PYK2 or by an extended KFL fragment (EGFP-KFL_700-841_) dissolved upon treatment with 5% of 1,6-HD for 30 s (**Fig. 4A**). Moreover, in live-cell imaging droplets fused into larger structures on a timescale of seconds (**Fig. 4B** and **Suppl. Video 1**). Fluorescence recovery after photobleaching (FRAP) revealed ≥70% recovery with a t_1⁄2_ of 9 ± 0.5 s (**Fig. 4C** and **Table 1**), indicating high mobility and exchange of molecules within the droplets^[37]^. These results showed that transfected EGFP-PYK2 forms puncta of which the number, size, and phase state are influenced by the PYK2 expression level.

**Fig. 4:**
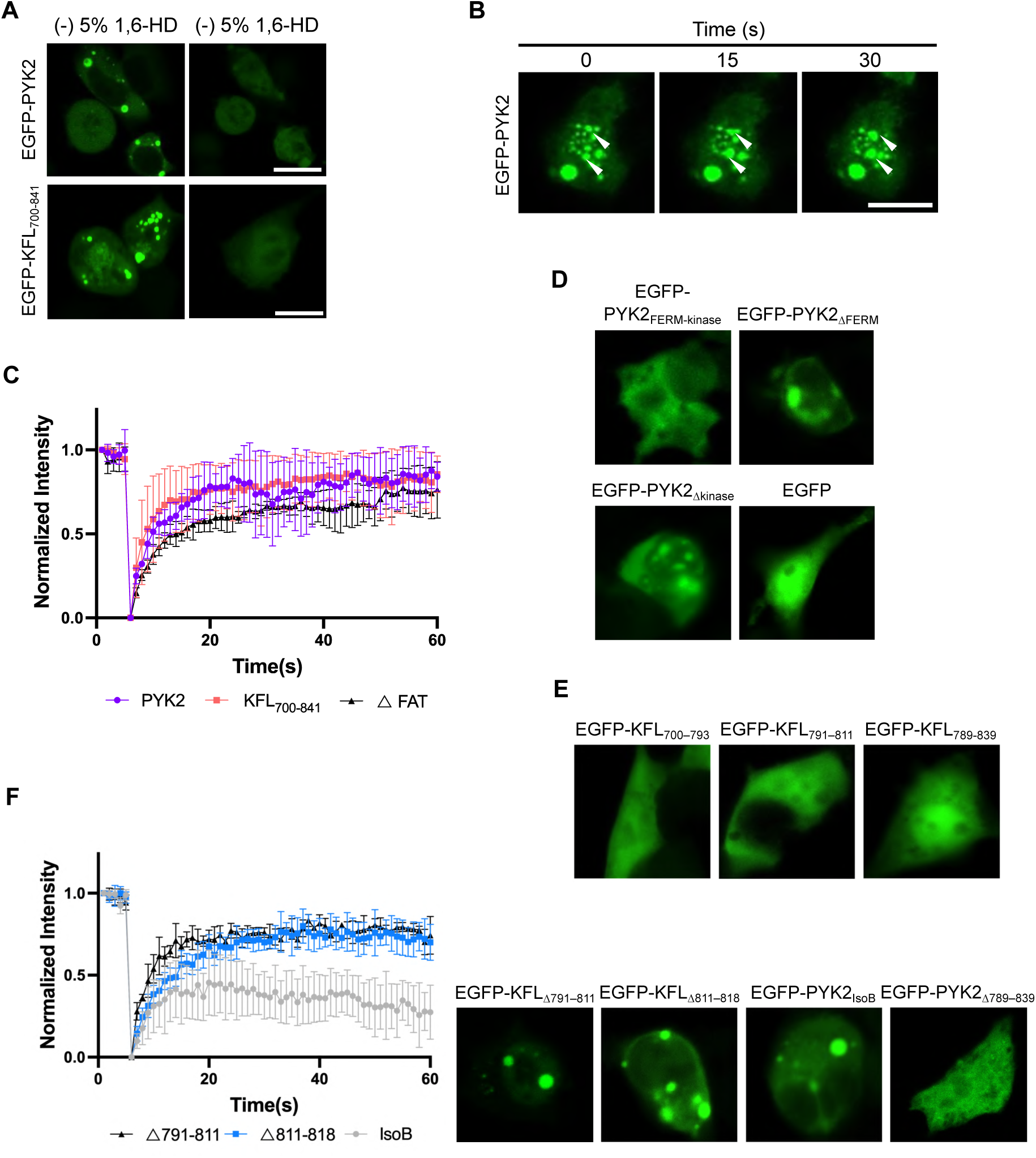
Distinct behavior of PYK2 constructs in cells reveals the critical role of the KFL helix in driving condensate formation. **A.** Fluorescence images of HeLa cells expressing EGFP-PYK2 (*top row*) or EGFP-KFL_700-841_ (*bottom row*) before and after 1,6-HD treatment (same field of view). Scale bar: 10 µm. Images were acquired with FLoid™ Cell Imaging Station. **B.** Time-lapse live-cell imaging of HeLa cells expressing EGFP-PYK2, showing droplet fusion events (white arrows) at 0, 15, and 30 seconds. Scale bar: 10 µm. Images were acquired with Leica Stellaris 8 FALCON confocal microscope equipped with a Zeiss 63x/1.4 oil objective. **C.** FRAP analysis of EGFP-PYK2, EGFP-KFL_700-841_, and EGFP-PYK2_ΔFAT_ puncta, showing fluorescence recovery curves (mean ± SEM, *n* = 3). **D.** Fluorescence microscopy images of HEK293T cells expressing domain-deletion variants: EGFP-PYK2_FERM-kinase_, EGFP-PYK2_ΔFERM_, EGFP-PYK2_Δkinase_, and EGFP alone (control). **E.** Fluorescence images of HEK293T cells transfected with EGFP-tagged KFL truncations: EGFP-KFL_700-793_, EGFP-KFL_791-811_, EGFP-KFL_789-839_. **F.** (*Left*) FRAP curves of EGFP-KFL_Δ791-811_; EGFP-KFL_Δ811-818_; EGFP-PYK2_IsoB_ droplets. (*Right*) Fluorescence microscopy images of HEK293T cells expressing EGFP-KFL_Δ791-811_; EGFP-KFL_Δ811-818_; EGFP-PYK2_IsoB_; EGFP-PYK2_Δ789-839_ plasmids. Images were acquired with FLoid™ Cell Imaging Station.

**Table 1:**
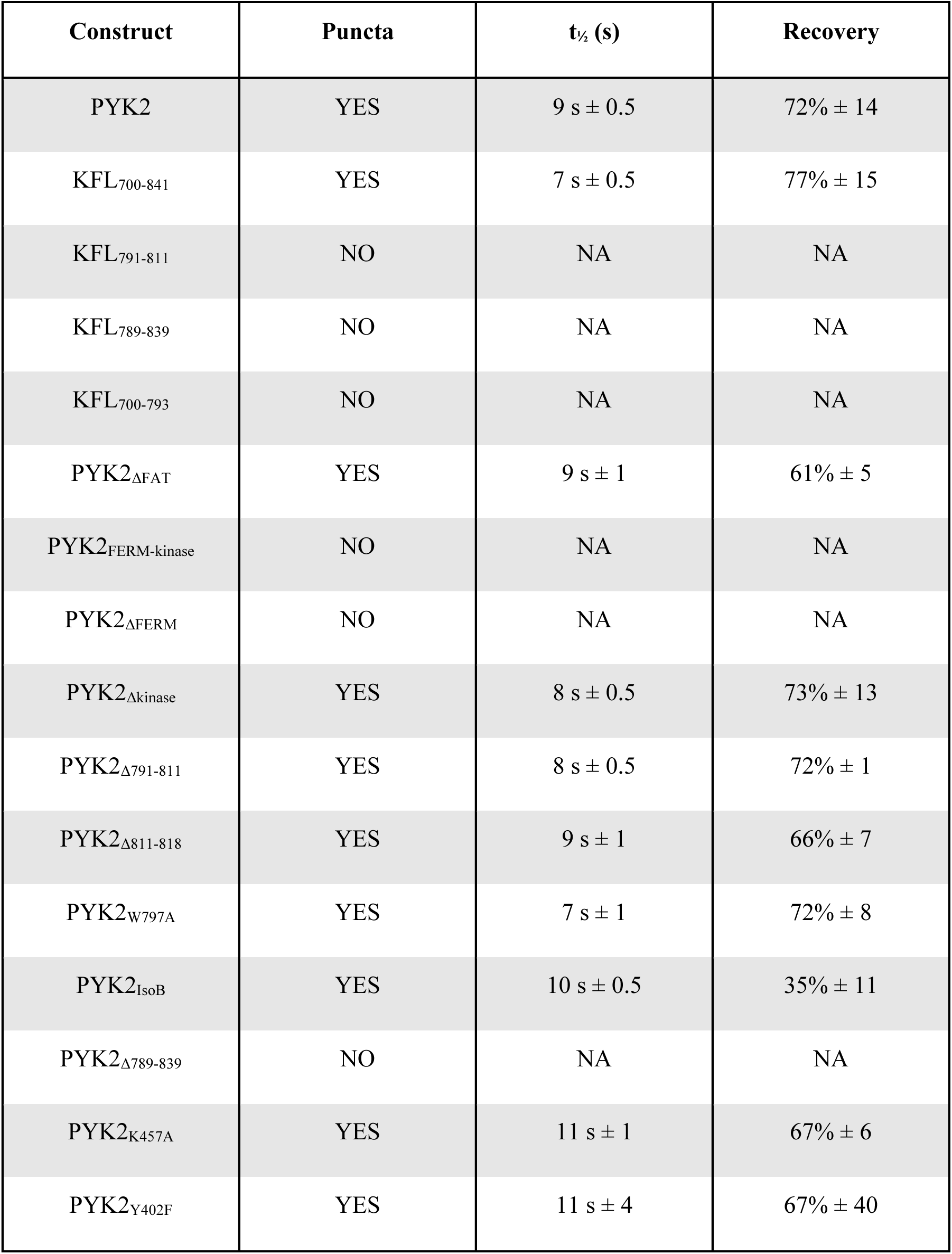
FRAP analysis and construct recovery dynamics. The recovery half-time is defined as the duration required for the bleached region to regain half of its final intensity. The recovery percentage is defined as the average intensity of the region after recovery relative to its initial intensity before bleaching.

| Construct | Puncta | $t_{1/2}$ (s) | Recovery |
| --- | --- | --- | --- |
| PYK2 | YES | $9 \text{ s} \pm 0.5$ | $72\% \pm 14$ |
| KFL<sub>700-841</sub> | YES | $7 \text{ s} \pm 0.5$ | $77\% \pm 15$ |
| KFL<sub>791-811</sub> | NO | NA | NA |
| KFL<sub>789-839</sub> | NO | NA | NA |
| KFL<sub>700-793</sub> | NO | NA | NA |
| PYK2<sub>ΔFAT</sub> | YES | $9 \text{ s} \pm 1$ | $61\% \pm 5$ |
| PYK2<sub>FERM-kinase</sub> | NO | NA | NA |
| PYK2<sub>ΔFERM</sub> | NO | NA | NA |
| PYK2<sub>Δkinase</sub> | YES | $8 \text{ s} \pm 0.5$ | $73\% \pm 13$ |
| PYK2<sub>Δ791-811</sub> | YES | $8 \text{ s} \pm 0.5$ | $72\% \pm 1$ |
| PYK2<sub>Δ811-818</sub> | YES | $9 \text{ s} \pm 1$ | $66\% \pm 7$ |
| PYK2<sub>W797A</sub> | YES | $7 \text{ s} \pm 1$ | $72\% \pm 8$ |
| PYK2<sub>IsoB</sub> | YES | $10 \text{ s} \pm 0.5$ | $35\% \pm 11$ |
| PYK2<sub>Δ789-839</sub> | NO | NA | NA |
| PYK2<sub>K457A</sub> | YES | $11 \text{ s} \pm 1$ | $67\% \pm 6$ |
| PYK2<sub>Y402F</sub> | YES | $11 \text{ s} \pm 4$ | $67\% \pm 40$ |

### The KFL drives PYK2 phase separation in cells

To dissect the drivers of cellular PYK2 phase separation, we transfected HEK293T cells with plasmids including different DNA constructs of PYK2 fused to EGFP (see **Fig. 1A** for PYK2 constructs nomenclature). When assessed after 24 hours, the KFL alone (EGFP-KFL_700-841_) and the PYK2 construct lacking the FAT domain (EGFP-PYK2_ΔFAT_) formed droplets that recovered from bleaching as fast as full-length EGFP-PYK2, 7 ± 0.5 s and 9 ± 1 s, respectively (**Fig. 4C** and **Table 1**). Conversely the FERM–kinase fragment (EGFP-PYK2_FERM-kinase_), lacking the KFL and FAT regions, formed static aggregates rather than dynamic droplets (**Fig. 4D** and **Table 1**). Deletion of the FERM domain (EGFP-PYK2_ΔFERM_) resulted in faint and diffuse puncta (**Fig. 4D** and **Table 1**). Disrupting the capability of the FERM domain to dimerise (through the mutation W273A) abolished formation of puncta (**Suppl. Fig. 3A**). Deletion of the kinase domain (EGFP-PYK2_Δkinase_) produced a construct that formed droplets that recovered well after photobleaching (≥70% recovery, t_½_ = 8 ± 0.5 s) at 16 hours, but rapidly transitioned to non-recovering aggregates at 24 hours (**Fig. 4D, Suppl. Fig. 3B** and **Table 1)**. EGFP alone failed to form droplets in HEK293T cells (**Fig. 4D**). These observations suggested that KFL is a driver of dynamic phase separation in PYK2, whereas the FERM-kinase domains promote aggregation. However, the collective action of KFL, FERM, and kinase determines the condensation dynamics in cells.

In the case of FAK, phase separation at concentrations as low as 40 nM was reported to be the result of multivalency arising from interactions of its FERM and FAT domains (i.e. FERM–FERM and FERM–FAT interactions), rather than from KFL interactions^[9,42]^. Conversely, in our experiments, PYK2_ΔFAT_ formed puncta, and PYK2 condensation relied on the KFL which is the least conserved region between FAK and PYK2 (**Fig. 1A, B**). Thus, the multivalency required for condensation is achieved differently in PYK2 and FAK.

### Determinants of phase separation within the KFL

To more finely dissect the molecular basis for KFL condensation, we transfected HEK293T cells with different PYK2 KFL constructs fused to EGFP (see **Fig. 1A** for KFL constructs nomenclature). The extended KFL fragment, containing also the PR2 region (EGFP-KFL_700-841_) formed puncta with fast FRAP (**Fig. 4A** and **Table 1**). EGFP-KFL_700-793_, encompassing only residues preceding the KFL helix (residues 791–811), failed to form droplets. However, neither the helix region alone (EGFP-KFL_791–811_) nor a larger fragment encompassing the helix (EGFP-KFL_789–839_) formed puncta (**Fig. 4E**).

Next, we probed the effect of deleting KFL residues in full-length PYK2. HEK293T cells transfected with PYK2 constructs that lacked either the KFL helix (EGFP-PYK2_Δ791–811_) or short segment thereafter (EGFP-PYK2_Δ811-818_) formed puncta that recovered after photobleaching (**Fig. 4F** and **Table 1**). In agreement, the mutation of a tryptophan within the helix to alanine (EGFP-PYK2_W797A_) did not abolish droplet formation and their FRAP (**Suppl. Fig. 3C** and **Table 1**). However, the deletion of a larger portion of the KFL region (EGFP-PYK2_Δ789–839_) completely abrogated the formation of liquid-like droplets, underscoring the essential role of this segment in driving LLPS of the protein (**Fig. 4F** and **Table 1**). We also tested the natural PYK2 isoform B (EGFP-PYK2_IsoB_) in which residues 739-780 are lacking. Although IsoB still formed puncta, their recovery after photobleaching was strongly impaired (**Fig. 4F** and **Table 1**). These results agreed with our NMR analysis indicating that the KFL helix is a key component for LLPS, however, droplet formation and dynamics result from fuzzy associations within the whole KFL.

### CaM is a client of PYK2 condensates

Calcium-sensing by PYK2 is mediated by the PYK2 KFL that forms a fuzzy association with CaM, which is strengthened fifteen-fold upon binding of Ca^2+^ to CaM^[30]^. Given that the KFL is driving PYK2 condensation, we asked whether CaM is recruited into PYK2 droplets. Indeed, i*n vitro*, we observed that CaM is recruited as a client into KFL_728-839_ droplets in the absence and presence of 5 mM CaCl_2_ (**Figure 5A**). CaM alone did not form droplets (**Suppl. Fig. 1B**). Next, we tested the effect of calcium on the PYK2 droplet formation in cells. We incubated HEK293T cells with 10 µM of a Calcium Ionophore or with 0.1% DMSO for 15 minutes and assessed the fluorescence intensity. Calcium-treated cells gained approximately 1.5 fold in EGFP-PYK2 fluorescence intensity (**Figure 5B**). This observation suggested that calcium-loaded CaM (Ca^2+^/CaM) facilitates the formation of PYK2 puncta.

**Fig. 5:**
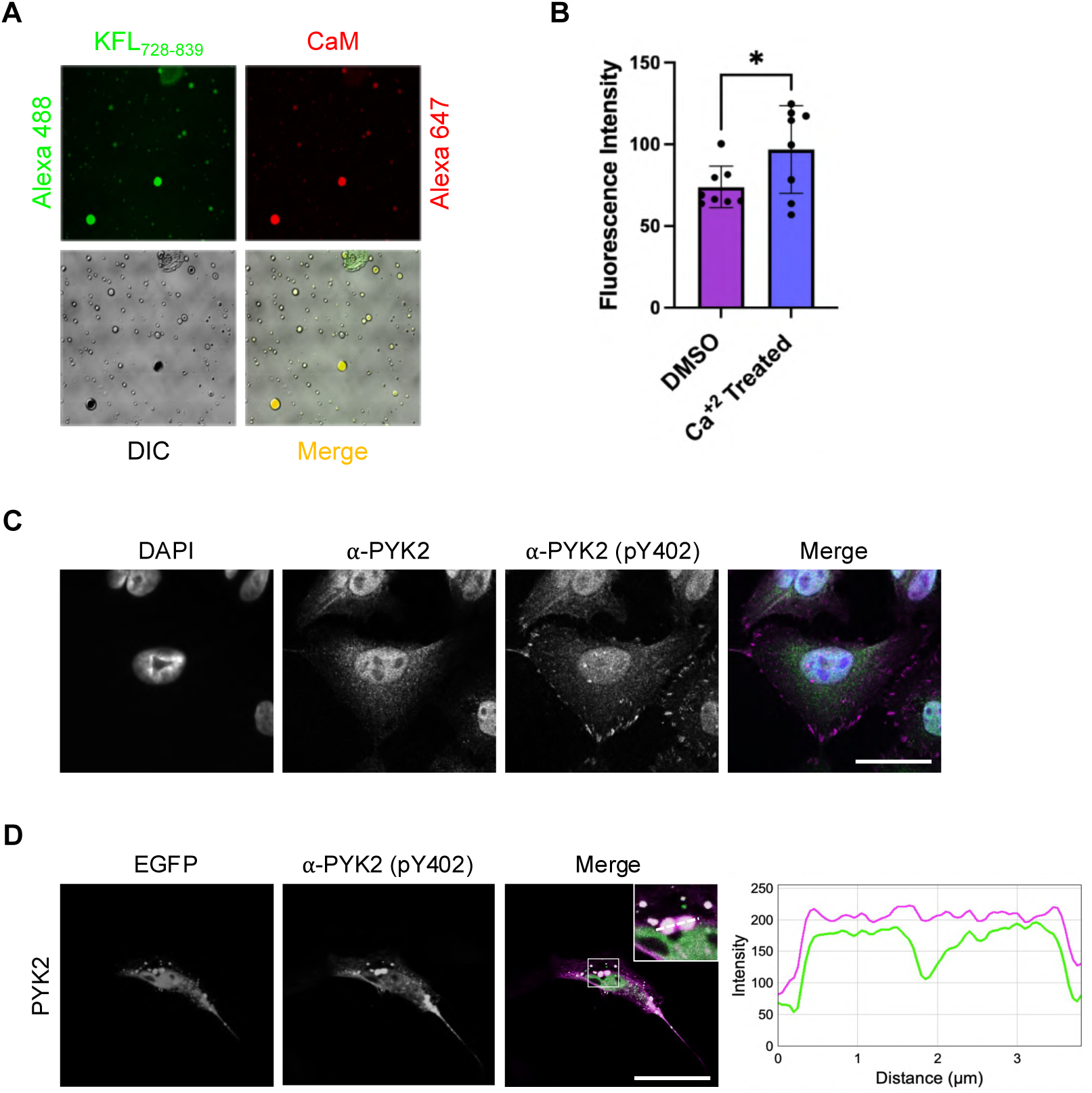
Calcium signaling and phosphorylation modulate PYK2 condensates. **A.** Widefield fluorescence and DIC microscopy images of droplets formed by Alexa 488-labelled KFL_728-839_ and Alexa 647-labelled CaM recruited as a client into KFL_728-839_ droplets with 200 mM NaCl. **B.** Fluorescence intensity measurements of EGFP-PYK2 in HEK293T cells treated with DMSO (control) or 10 µM Calcium Ionophore (Ca^2+^ treated). *p < 0.05 **C.** Co-immunostaining of HeLa cells with antibodies against total PYK2 (green), phosphorylated PYK2 (pY402, magenta), and DAPI (blue), shown as individual and merged channels. **D.** Immunostaining of HeLa cells expressing EGFP-PYK2 with anti-pY402 antibody (green and magenta in merged images), with corresponding intensity profile plots along the indicated line. Scale bar: 10 µm. Images were acquired using a Leica Stellaris 8 FALCON confocal microscope with a Zeiss 63x/1.4 oil objective. CaM: calmodulin.

### Many, but not all, cytoplasmic PYK2 puncta contain phosphorylated Y402

Local enrichment and self-association is essential for the autophosphorylation *in trans* of residue Y397 of FAK at IACs^[42,43]^. Given that the phosphorylation *in trans* is likely conserved in PYK2 as Y402^[22,44]^, and that Ca^2+^/CaM enhances KFL dimerisation *in vitro*^[30]^, we tested whether condensates contained PYK2 phosphorylated on Y402. Confocal imaging of HeLa cells using a total and a pY402-specific PYK2 antibody showed that PYK2 molecules localising at IACs were highly autophosphorylated, in line with a model where local enrichment of FAK and PYK2 leads to their self-association and autophosophorylation *in trans*^[42]^. Many, but not all, cytoplasmic PYK2 puncta were also marked by the pY402-specific antibody (**Fig. 5C**). Immunostaining of EGFP-PYK2 transfected HeLa cells using this PYK2 pY402 antibody showed that most of these PYK2 puncta were also enriched in phospho-Y402 (**Fig. 5D**). Confocal microscopy showed that EGFP-PYK2_Y402F_ and kinase-dead EGFP-PYK2_K457A_ mutants formed droplets that recovered after photobleaching (**Table 1**), but also aggregates (**Suppl. Fig. 3D**). These observations suggested that condensation promotes autophosphorylation of PYK2 by presenting an environment where PYK2 is enriched, rather than that autophosphorylation promotes PYK2 condensation.

### PYK2 condensates recruit paxillin, weakening cell attachment

We next investigated the cellular effects of PYK2 condensates. PYK2 interacts with many proteins, including paxillin^[45,46]^. Both PYK2 and FAK use their FAT domains to associate with the leucine-aspartic acid (LD) motifs of paxillin^[17,32,47,48,49,50]^. Paxillin localizes to the nucleus, cytoplasm, and membrane-proximal regions, with the latter being critical for IAC formation through paxillin-facilitated phase separation^[9,51]^. Interestingly, when immunostaining paxillin in HeLa cells overexpressing EGFP-PYK2, we observed that paxillin mainly colocalizes with perinuclear EGFP-PYK2 puncta (**Fig. 6A**). In contrast, untransfected cells with lower endogenous PYK2 levels showed paxillin primarily at IACs and, to a lesser extent, within the nucleus (**Fig. 6A, B**).

**Fig. 6:**
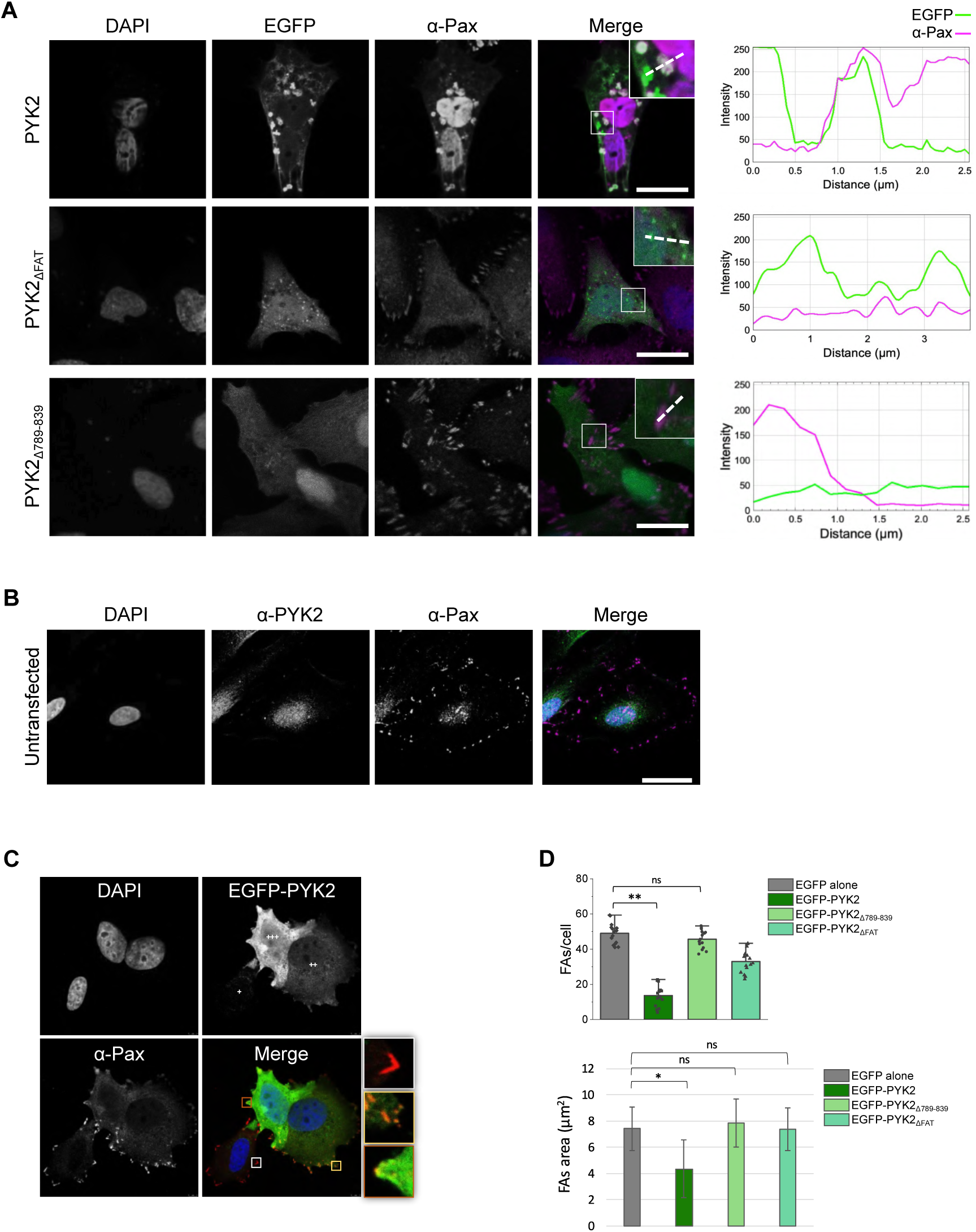
PYK2 condensates modulates FAs disassembly by interacting with paxillin in cells. **A.** Fluorescence images of HeLa cells expressing EGFP-PYK2 (*top row*), EGFP-PYK2_ΔFAT_ (*middle row*) or EGFP-PYK2_Δ789-839_ (*bottom row*) in green, immunostained with anti-paxillin (magenta) and counterstained with DAPI (blue). Intensity profile plots on the right show signal distribution along the indicated line for PYK2 variants and paxillin. **B.** Co-immunostaining of endogenous PYK2 (green), paxillin (magenta), and DAPI (blue) in HeLa cells. **C.** MCF-7 cells showing various degrees of EGFP-PYK2 expression (+, ++, +++) in green and stained with anti-paxillin antibody (magenta). Color-matched zoomed views of individual IACs are shown on the right. **D.** (*Top*) Quantification of FA number per cell in cells untransfected or transfected with EGFP-PYK2, EGFP-PYK2_Δ789-839_ or EGFP-PYK2_ΔFAT_. (Bottom) Bar plot of the FA area per cell under the same conditions (mean ± SEM, *n* = 30). *p < 0.05, **p < 0.01 compared to EGFP alone (control). Pax: paxillin.

To investigate whether the FAT-LD interaction is necessary for paxillin recruitment to PYK2 condensates, we transfected cells with EGFP-PYK2_ΔFAT_, a mutant retaining the ability to form droplets (**Fig. 4C, 6A and Table 1**). EGFP-PYK2_ΔFAT_ condensates failed to recruit paxillin, which remained localised at FAs (**Fig. 6A**), emphasising the critical role of the FAT domain in mediating the binding between PYK2 and paxillin, and the ability of PYK2 to undergo phase separation independently of this interaction. Remarkably, paxillin exclusively localised at IACs in cells transfected with EGFP-PYK2_Δ789–839_, a mutant that retains the FAT domain but is incapable of phase separation (**Fig. 4F** and **6A**). Thus, both the presence of the FAT domain and the ability of PYK2 to phase separate are required to recruit paxillin away from IACs, consequently weakening cellular attachment (**Fig 6A** and **7A**).

**Fig. 7:**
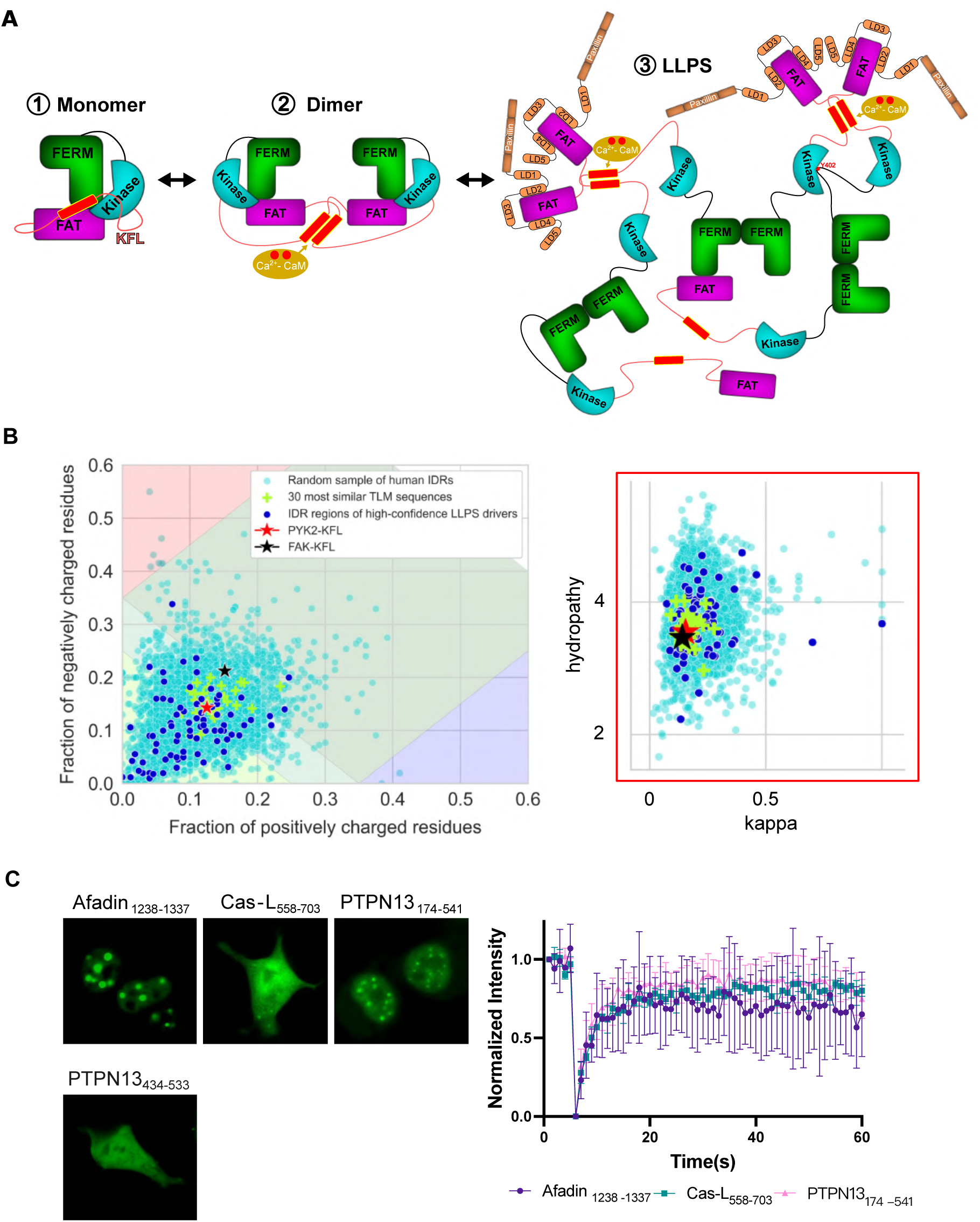
PYK2 LLPS expands the functional condensates landscape. **A.** Schematic model of PYK2 phase separation via dimerization and interaction with Ca^2+^/CaM and paxillin, highlighting its role in condensate formation and function. **B.** (*Left)* Diagramme of states showing the distribution of positive and negative charges for a sample of 5,000 human IDRs, the 30 most similar IDRs to the PYK2 KFL sequence, and fragments from 83 high-confidence LLPS drivers that satisfied the same criteria as for the human IDRs (see Methods). The regions of the diagramme of states are as follows: yellow, weak polyampholytes that assume globular and "tadpole" conformations; light green, Janus sequences that can be collapsed or expanded depending on their context; dark green, strong polyampholytes that form coils, hairpins and chimaeras; red, negatively charged strong polyelectrolytes that behave as swollen coils; blue, positively charged strong polyelectrolytes that also behave as swollen coils. (*Right)* mean hydropathy vs kappa value. kappa is a parameter to describe the degree of mixing of negatively and positively-charged amino acids. Values closer to 1 indicate a segregation of positive and negative charges, and values closer to 0 indicate an even mixing of positive and negative charges. **C.** (*Left*) Fluorescence images of HEK293T cells expressing EGFP-Afadin_1238-1337_, EGFP-PTPN13_174-541,_, EGFP-Cas-L_558-703_, EGFP-PTPN13_434-533_. Images were acquired with FLoid™ Cell Imaging Station. (*Right*) FRAP curves of the constructs forming puncta on the left (mean ± SEM, *n* = 3).

Consistent with these findings, MCF-7 breast cancer cells transfected with EGFP-PYK2 display an inverse correlation between EGFP-PYK2 expression levels and the presence of paxillin-positive FAs (**Fig. 6C**). Higher expression levels of EGFP-PYK2 correlated with increased colocalization of paxillin in PYK2-containing puncta (**Fig. 6A, C**). At moderate expression levels, however, paxillin colocalized with only a subset of these puncta, suggesting structural or functional heterogeneity among PYK2 condensates (**Fig. 6A, C**). Functionally, overexpression of full-length PYK2 results in a marked decrease in both the number and area of FAs compared to untransfected cells (**Fig. 6D**). These cells also exhibit a distinct morphology, appearing partially lifted from or loosely attached to the substrate (**Suppl. Fig. 3E**). These data support that the redistribution of paxillin into EGFP-PYK2 condensates depends on PYK2 expression levels, and results in decreased cell attachment and reduced FAs (**Fig. 6C, D** and **Suppl. Fig. 3E**).

In line with this model, TIRF showed that the population of EGFP-PYK2 that is localised at HeLa IACs is getting depleted as these IACs are disassembled at the cell trailing edge (**Suppl. Video 2**). Given this observation, a mechanism where budding PYK2 droplets drag paxillin away from IACs would be appealing^[16]^. However, we did not observe PYK2 droplet budding from IACs under either medium (20 µg/mL) or high (50 µg/mL) fibronectin conditions (**Suppl. Video 2 and 3)**.

### The KFL extends the range of LLPS drivers

Having observed that KFL LLPS was unaffected by phosphorylation and that the KFL lacked apparent multivalency *in vitro*, we investigated whether the KFL is an atypical driver of phase separation. To this end, we compared the PYK2 KFL sequence against a curated set of 89 high-confidence LLPS drivers^[52]^. We also constructed a database of all human intrinsically disordered regions (IDRs). This IDR database was constructed from AlphaFold-predicted human proteome structures^[53]^, selecting regions with: (a) a low proportion of residues with pLDDT > 80 (to permit short secondary structures), (b) a minimum length of 50 residues, and (c) varying sequence lengths (see **Methods**). Combining 18 combinations of maximum percentage of pLDDT>80 and sequence window length we compiled a human IDR database with 8,557,526 unique sequences. We then used ESM-2, a transformer protein language model^[54]^, to implement a sequence embedding–based selection criterion to rank the sequences in our IDR database according to their similarity to the PYK2 KFL sequence. This approach was necessary due to the lack of proteins with significant sequence identity to PYK2 KFL beyond the ∼20% identical FAK KFL. We then plotted the high-confidence LLPS drivers, the 30 IDR sequences most similar to PYK2 KFL and 5,000 randomly selected IDR sequences using physicochemical criteria relevant to IDRs(**Fig. 7B** and **Suppl. Fig. 4**)^[55]^.

In the diagram of states—which plots the fraction of positively versus negatively charged residues—the PYK2 KFL sequence falls within the region associated with “Janus” sequences, characterized by a near-neutral net charge (**Fig. 7B**). These Janus sequences are notable for their ability to adopt either collapsed or extended conformations depending on the physicochemical environment. In contrast, most known LLPS drivers clustered in a less charged region associated with more compact conformations, often referred to as “globules and tadpoles.” The randomly selected set of 5,000 IDRs spans the full range of the diagram, with only a few sequences occupying the highly charged “swollen coils” region. To further contextualize KFL’s position, we extended the analysis using ten additional physicochemical criteria—such as hydropathy and charge patterning (e.g., the kappa score)—resulting in 55 two-dimensional plots (**Suppl. Fig. 4**). All these representations recapitulated the observations seen in the diagram of states: The PYK2 KFL sequence consistently occupied a central, average position within the human IDR landscape, closely surrounded by sequences with the most similar ESM-2 embeddings; conversely known LLPS drivers were more dispersed and generally more distant from KFL. This analysis supported that the Janus-class KFL is atypical among known LLPS sequences, whereas many human IDRs share similar sequence features, and hence potentially similar phase separation capabilities.

A gene ontology (GO) analysis of 150 genes that were ranked close to KFL by ESM-2 showed an enrichment in cytoplasmic proteins involved in cytoskeletal organisation (in particular actin and microtubules) (see **Methods** and **Suppl. Fig. 5**). Other enriched terms were adhesions (FAs, synaptic structures) and intracellular signal transduction. These GO terms were closely mirroring the functional profile of PYK2. To experimentally assess whether these KFL-like IDRs could drive phase separation, we selected three sequences from proteins identified as highly similar to the KFL by ESM-2: Afadin, PTPN13 and Cas-L. Following their transfection into HEK293T cells as EGFP fusion constructs, we found that EGFP-Afadin_1238-1337_, EGFP-PTPN13_174-541_, and EGFP-Cas-L_558-703_, formed droplets that recovered after photobleaching (**Fig. 7C**). The shorter EGFP-PTPN13_434-533_ construct did not form droplets under the same conditions, supporting that droplet formation is not inherent to our experimental design (**Fig. 7C**). Afadin, PTPN13, and Cas-L function as key regulators of cell adhesion dynamics and cytoskeletal remodeling, integrating signaling pathways that control cell polarity, migration, and tissue organization^[56,57,58]^. While the physiological relevance of LLPS in these proteins remains to be established, our findings suggest that Janus-type IDRs may represent an underappreciated class of homotypic LLPS drivers, particularly enriched at the intersection of cytoskeletal dynamics, adhesion, and intracellular signaling.

## DISCUSSION

This study provides the first detailed mechanistic and cellular characterization of phase separation by PYK2, revealing a condensation mechanism that diverges from its close paralogue, FAK. Through mutational analysis, we identified the KFL region as the primary driver of dynamic LLPS in PYK2, while the FERM and kinase domains promote aggregation. The balance and interplay between these domains determines the overall condensation behavior of PYK2. This contrasts with FAK: although one group observed that the isolated FAK KFL sequence forms droplets *in vitro* under high concentrations of protein and crowding agents^[59]^, condensation of full-length FAK is driven by multivalent associations between its FERM and FAT domains^[9]^. Indeed, KFL is the region that differs most between both paralogues and it is also the site of the PYK2-specific CaM interaction^[30]^. Thus, differentiation within the KFL emerges as a key mechanism that allowed PYK2 to adapt the ancestral FAK framework for different biological roles.

Using NMR spectroscopy, we found that the LLPS transition in PYK2 KFL involves only subtle CSPs, particularly within the helical region previously implicated in dimerization and CaM binding^[30]^. However, this region alone is insufficient to drive LLPS, and its disruption *via* mutations or short deletions did not fully abolish condensation. These data are consistent with condensation resulting from many dynamic, distributed, low-affinity interactions across the disordered protein sequence—a concept recognized in the LLPS field^[60]^. Surprisingly, *in vitro* PYK2 KFL condensation was insensitive to phosphorylation. This contrasts with many IDR-driven condensates where phosphorylation modulates phase behavior. The lack of effect in PYK2 is likely due to the spatial separation of phosphorylation sites from the key interaction regions, as confirmed by NMR and mutational analysis.

We demonstrate that endogenous PYK2 forms mostly cytoplasmic puncta across various cell types. These puncta are distinct from endosomes and sensitive to salt and 1,6-HD, indicating their liquid-like nature. This pattern contrasts with FAK, which forms membrane-associated condensates through cooperation with paxillin and other FAs proteins to promote adhesion^[9]^. It is unclear why PYK2 condensates nucleate in the cytoplasm, rather than at sites of adhesions akin to FAK. Possible explanations include the additional recruitment of FAK to IACs by other factors that do not interact with PYK2 (e.g. talin^[34]^), the retention of PYK2 in the cytoplasm by specific partners. Another possibility is represented by p130Cas, which was observed to form condensates that bud from FAs when cells were plated at high fibronectin concentrations^[16]^. Although we detected residual PYK2 at IACs by super-resolution microscopy, we failed to observe droplet formation from IACs under the tested conditions.

Functionally, we show that Ca^2+^/CaM acts as a client of PYK2 condensates *in vitro* and promotes puncta formation in cells, suggesting a role for calcium signaling in modulating PYK2 phase behavior. Furthermore, we find that most condensates are enriched in Y402-phosphorylated PYK2, representative of PYK2 as a hub for kinase activity and signalling scaffold. This observation supports a model in which condensation facilitates autophosphorylation by increasing local concentration and promoting trans-activation^[61]^. This mechanism is conceptually similar to FAK activation at IACs, where local enrichment of paxillin and other factors drives self-association and phosphorylation *in trans*^[42,43]^.

Importantly, our work highlights a substantial dependency of the number, size, phase state, and biological effect of PYK2 condensates with PYK2 expression levels: at low/endogenous expression levels PYK2 produces small cytoplasmic, perinuclear, and nuclear puncta, some of which are enriched in pY402-PYK2, suggesting they function as dispersed signalling hubs. Moderate overexpression enhanced the presence of PYK2 at IACs, but also promoted the recruitment of paxillin into cytoplasmic PYK2 puncta. Both the turnover of PYK2-containing IACs and cytoplasmic sequestration of paxillin required the KFL region that is essential for PYK2 phase-separation. Further increases of PYK2 expression led to the formation of large, predominantly periplasmic condensates that fully sequester paxillin away from FAs, resulting in cell detachment. Under these conditions PYK2 droplets also captured actin, disrupting cytoskeletal architecture (**Suppl. Fig. 3F)**. At even higher concentrations PYK2 forms large static aggregates.

This concentration–localization–function relationship mirrors recent findings in other phase-separating systems^[62,63,64,65]^, and underscores the importance of tightly regulating PYK2 expression in cells. Moreover, our findings caution that non-physiological overexpression of PYK2 can generate aberrant condensates with potentially misleading or deleterious effects. Given that PYK2 is overexpressed in several aggressive malignancies^[26,27,66,67]^, our findings raise the possibility that aberrant condensates may contribute to adhesion defects by pathologically weakening cell–matrix interactions. We did not observe paxillin sequestration by endogenous PYK2 in our cell lines and under the experimental conditions used. However, potential pathogenic effects of aberrant PYK2 condensates, possibly as a result of stimuli-induced enhanced PYK2 expression (e.g. through EGF stimulation or following calcium influx), remain a possibility and require further investigations.

Previous findings suggested, collectively, that the local enrichment of FAK at IACs leads to its FERM-mediated self-association, which is required for subsequent *trans*-autophosphorylation, and hence catalytic activation^[9,42,68,43]^. Jointly with our previous investigations^[30]^, our data suggest that for PYK2, this mechanism for *trans*-autophosphorylation is also based on local enrichment. However, this enrichment occurs in cytoplasmic condensates, where the KFL is freed from its inhibitory association *in cis* with the FERM-kinase domains, to form dimers and higher-order fuzzy self-associations. Hence, our data support that PYK2 condensation is mechanistically linked to its catalytic activity. It remains to be determined whether the PYK2 condensates nucleate stochastically in the cytoplasm at certain cellular PYK2 concentrations, and/or are promoted by other factors. For example, our observations support that calcium-bound CaM serves as a client for PYK2 puncta and further enhances self-association. Moreover, although phosphorylation of the KFL by other kinases did not alter its LLPS propensity *in vitro*, in cells the phosphorylation of the KFL may weaken its inhibitory association with the FERM-kinase fragment. In addition, the exposure of its PR2 and PR3 interaction sites may alter PYK2 condensates by recruiting additional molecules as clients. The nature and role of additional partners in condensate function and the dynamics of their transition to aggregates remains to be explored.

Akin to PYK2, ligand-induced dimerization was also shown to promote LLPS in some receptor tyrosine kinases, such as FGFR2 and EGFR^[69,70,11]^. Yet, in these cases, LLPS occurred through a different mechanism, namely by creating multiple phosphotyrosine-based protein interaction motifs that trigger heterotypic condensates through specific multivalent associations with multivalent scaffolding phosphotyrosine reader domains^[11]^. Mechanistically, LLPS autoregulation by PYK2 displays some similarity to the one identified for protein kinase A (PKA). cAMP-promoted dissociation of the catalytic and regulatory subunits of PKA leads to the exposure of an otherwise hidden flexible region that allows PKA to associate with heterotypic condensates^[71]^. However, the triggers and functional repercussions of LLPS driver exposure differ markedly also between PKA and PYK2. Hence, PYK2 expands the range of molecular mechanisms used for controlling phase separation.

Finally, we show that the PYK2 KFL sequence belongs to the “Janus” class sequences, characterized by a near-neutral net charge and notable for their ability to adopt either collapsed or extended conformations depending on the physicochemical environment^[55,72,73]^. Our analysis supported that the Janus-class drivers are atypical among known LLPS sequences. However, using the transformer-based protein language model ESM-2, we identified a broader class of IDRs with sequence features similar to the Janus-type KFL, suggesting that these IDRs, found in proteins involved in adhesion and cytoskeletal regulation, also exhibit phase-separating potential in cells. In support, We confirmed the phase-separating potential in cells for sequences derived from Afadin, PTPN13, and Cas-L. Therefore, we propose that Janus-type IDRs may represent an underappreciated class of homotypic LLPS drivers, particularly enriched at the interface of cytoskeletal dynamics, adhesion, and signaling.

In conclusion, our study reveals a novel mechanism linking kinase activation to phase separation, highlights functional divergence between closely related kinases, and expands the landscape of LLPS drivers into Janus-class sequences. These findings also underscore the importance of protein dosage in modulating condensate behavior and cellular effects, with potential implications for understanding signaling dynamics and disease-associated dysregulation.

## MATERIALS AND METHODS

### Recombinant cloning and protein production

The recombinant cloning was performed using restriction digestion with BamHI, XhoI and ligated in pET32-a (+) vector with an N-terminal HisTag for FAK (FAK-KFL_764–845_, FAK-KFL_776–841_, FAK-KFL_804–832_), PYK2 (KFL_728–839_, KFL_761–839_, KFL_789–839_) and CaM (CaM_1-148_). DNA fragments from *Homo sapiens* were amplified using the oligonucleotide primers described in *Momin et, al* ^[30]^. The proteins were expressed and purified as described in *Momin et, al*^[30,37]^ with minimal differences. Briefly, the cloned vectors were expressed in *Escherichia coli* cells induced with 0.5 mM IPTG. 10 mM imidazole was added to the wash buffer for the Ni-NTA column affinity purification, and proteins were eluted at a gradient elution using ÄKTA Pure system. Eluted proteins were subjected to a Superdex-75 size-exclusion column (Cytiva) in a SEC buffer containing 20 mM HEPES pH 7.5, 200 mM NaCl, and 1 mM TCEP. The purity for all proteins was measured using SDS-PAGE gels and all proteins were stored at -80°C in the SEC buffer.

The His-tagged EGFP-PYK2 kinase domain protein was expressed in CHO mammalian cells using the ExpiCHO™ Expression System Kit (ThermoFisher, CAT: A29133). In a 250 mL flask, ExpiCHO-S™ Cells were diluted to a final density of 6 x 10^6^ viable cells/mL with fresh ExpiCHO™ Expression Medium, pre-warmed to 37°C. ExpiFectamine™ CHO/plasmid DNA complexes were prepared using cold reagents (4 °C): 40 µg of plasmid DNA was diluted in a cold OptiPRO™ medium to a final volume of 2 mL, and 160 µL ExpiFectamine™ CHO Reagent was diluted in OptiPRO™ medium to a final volume of 2 mL. The ExpiFectamine™ CHO/plasmid DNA complexes were incubated at room temperature for 5 minutes, and then the solution was slowly transferred to the shaker flask with ExpiCHO-S™ Cells, swirling the flask gently during the addition. The cells were then incubated in a 37°C incubator with a humidified atmosphere of 8% CO2 in the air on an orbital shaker (120 RPM). On the day after transfection, 300 µL ExpiFectamine™ CHO Enhancer and 12 mL ExpiCHO™ Feed were added to the cultured cells, and the flask was returned to the 37°C incubator with a humidified atmosphere of 8% CO2 with shaking. The cells were harvested 10 days post-transfection by centrifugation in a benchtop centrifuge for 20 minutes at 4000 x g and 4◦C. The cells were lysed using lysis buffer (50 mM Tris-HCL (pH 7.4), 150 mM NaCl, 1% Triton, 1X RIPA buffer, 1 mM PMSF, 2 mM DTT, 1X Protease inhibitor EDTA-Free, Benzonase 25 U/ml). Followed by two-step FPLC purification using affinity (HisTrap HP, 5 ml) with binding buffer consisted of 25 mM Tris (pH 7.4), 500 mM NaCl, 10 mM Imidazole, 10% Glycerol, and 2 mM DTT, while the elution buffer consisted of 25 mM Tris (pH 7.4), 500 mM NaCl, 500 mM Imidazole, 10% Glycerol and 2 mM DTT. Size Exclusion Chromatography (SEC) was performed in Superdex™ 75 Increase 10/300 GL column. The elution buffer consisted of 20 mM HEPES, 300 mM NaCl, 10% Glycerol, and 2 mM DTT (pH 8.0).

### Nuclear Magnetic Resonance (NMR) spectroscopy measurements and analysis

NMR data were acquired on 700 MHz Bruker Avance NEO spectrometers equipped with 5 mm direct-detected (N/C/H-D) TXO cryogenic probe. Measurements were carried out at 37°C (310 K) on 250 μM KFL_728-839_ in 20 mM HEPES, 200 mM NaCl, 1 mM TCEP, pH 6.5 and pH 7.5), with 1%/99% (*v*/*v*) D_2_O/H_2_O. NoLLPS sample was measured by adding 5% 1-6, HD to the buffer and LLPS sample was measured after 24 h when it forms complete LLPS. NMR data processing and interpretation were performed using Topspin ver. 4.0.5, SPARKY (https://www.cgl.ucsf.edu/home/sparky/) and CARA (http://cara.nmr.ch/) software.

The assignments for backbone and side-chain ^1^H, ^13^C, ^15^N resonances were previously published. Additionally, ^13^C-detected experiment, 2D (H)CαCO (c_hcaco_ia3d, 16 scans), (H)CβCαCO (c_hcbcaco_ia3d, 32 scans), and (H)CACON (c_hcacon_ia2d, 32 scans), were recorded in order to enhance the sensitivity. (H)CACON spectra were utilised to obtain peak intensities and Linewidth (Iw1, Iw2) of LLPS divided by NoLLPS of PYK2 KFL_728-839_. CSP for ^13^C and ^15^N was calculated as follow

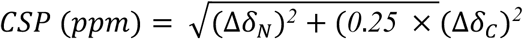

The median and IQR values were calculated (equation IQR = 1.5 * (Q_3_ – Q_1_, where Q_1_ is the first quartile, and Q_3_ is the third quartile). The line cutoff value was obtained by the addition of the median to IQR.

### ADP-Glo™ Kinase Assay

The ADP-Glo™ Kinase Assay Kit (Promega) was used to confirm the enzymatic activity of Src and MAPK kinases. Both enzymes were diluted in 1X kinase reaction buffer containing 40 mM Tris-HCl (pH 7.5), 20 mM MgCl₂, 2 mM MnCl₂, 0.1 mg/mL BSA, 2 mM DTT, and 100 µM ultra-pure ATP. Kinases were tested at two concentrations: 500 nM and 2500 nM. As a substrate, a recombinant PYK2 Kinase-FAT linker (KFL_728–839_) was used at a final concentration of 2 µM. Reactions were set up in a 384-well plate with a 5 µL reaction volume. After 2 hours of kinase reaction, 5 μL of ADP-Glo™ Reagent was added and incubated for 40 minutes to terminate the reaction. Subsequently, 10 µL of Kinase Detection Reagent was added to convert ADP to ATP and initiate the bioluminescent reaction. Luminescence was measured using a Tecan Infinite® M1000 microplate reader.

### Mass-spectrometry analysis

The phosphorylation reaction was kept at 50 µL containing 10 µM of KFL_728-839_ with 5 µM of MAPK or Src kinase in 50 mM Tris HCl (pH 7.5), 200 mM NaCl, 5 mM MgCl_2_, and 1 mM ATP. Mixed proteins were incubated for 1 hour and then run on SDS PAGE gel. The KFL_728-839_ bands were cut and underwent in-gel trypsin digestion, desalting, and cleanup. Mass-spectrometry analysis was performed as previously described in *Momin et al 2019*^[37]^, with the difference in data analysis performed using Mascot peptide mass fingerprinting software to identify modified residues in the KFL_728-839_.

### Mammalian cell culture

Human fibroblast (HDFa, PCS-201-012), HEK293T (human embryonic kidney, CRL-3216), HeLa (cervical cancer, CCL-2) and MCF-7 (non-invasive breast cancer, HTB-22) cells were cultured in high-glucose DMEM supplemented with 10% FBS (VWR, 10082147) and Penicillin-Streptomycin (Gibco, 15070063) at 37°C in a humidified atmosphere containing 5% CO₂. MDA-MB-231 (invasive breast cancer, HTB-26) cells were grown in L-15 medium supplemented with 10% FBS (VWR, 10082147) and Penicillin-Streptomycin (Gibco, 15070063), and cultured at 37°C in 5% CO_2_. Adherent cells were passaged every 3–4 days upon reaching 70–80% confluency. Cells were detached using 0.25% Trypsin-EDTA (ThermoFisher, 25200056) for 2-3 minutes, collected, resuspended in fresh complete medium, and reseeded. THP-1 (human acute monocytic leukaemia, TIB-202) cells were cultured in suspension in RPMI-1640 medium supplemented with 10% FBS and 0.05 mM 2-mercaptoethanol at 37°C in 5% CO_2_. Cells were subcultured every 3-4 days when reaching the concentration of 8 x 10^5^ cells/mL.

For the experiments including Ca^2+^, HEK293T cells were treated with 10 µM of Calcium Ionophore (Sigma, A23187) or 0.1% DMSO (vehicle control) and then incubated at 37℃ for 15 minutes. Following the incubation period, fluorescence intensity was measured in both conditions.

For the 1,6-HD treatments, live cell imaging was performed on an inverted Zeiss LSM880 confocal microscope. HEK293T cells transfected with various EGFP-PYK2 fusion constructs were cultured in a 35 mm glass-bottom dish until approximately 70% confluency and then imaged using the 488 nm laser. The working solution of 1,6-HD (Sigma-Aldrich, 240117) at a concentration of 5% in phenol-red free medium was freshly prepared. Following the initiation of image acquisition, pre-warmed hexanediol was added to the dish. The time of hexanediol addition was designated as time 0.

### Bacterial transformation and plasmid purification

One ShotTM Mach1^TM^ T1 Phage-Resistant Chemically Competent E. coli (ThermoFisher, C862003) cells were utilised following the transformation procedure outlined by the manufacturer’s instructions. Subsequently, plasmid purification was carried out using the QIAfilter EndoFree Plasmid Kit (QIAGEN, 12362) following the manufacturer’s instructions.

### Transient transfection

Starting with the manufacturer’s recommended amount of DNA equivalent to approximately 250 ng/cm², we systematically adjusted the DNA concentration while maintaining constant cell seeding density, incubation time and lipids ratios (**Suppl. Fig. 2D**). Cells were transfected with Lipofectamine™ 3000 (ThermoFisher, L3000001) following the manufacturer’s instructions. In brief, 24 hours after seeding on 35 mm glass-bottom dishes (Willco, 10810-054) or 6-well plates (Corning, CLS 3516), cells at 80–90% confluency were transfected. The complete medium was replaced with a transfection mix containing FBS-depleted medium, plasmid DNA (900 ng), Lipofectamine™ 3000 reagent, and P3000™ reagent at a 1:2:1 ratio. After incubation for 6 hours at 37°C in 5% CO₂, the medium was replaced with a complete medium. Transfection efficiency was assessed using the FLoid™ Cell Imaging Station 24-72 hours post-transfection.

### Immunofluorescence

Approximately 1 x 10^5^ cells were seeded in 35 mm glass-bottom dishes. After 24 hours, the medium was removed and cells were washed with PBS. Cells were fixed with freshly prepared PFA 4% (ThermoFisher, 28908) for 15 minutes, permeabilized with 0.2% Triton X-100 in PBS for 30 minutes, and washed three times with PBS. Blocking was performed for 1 hour in BSA 3% in PBS. Cells were then incubated overnight at 4°C with primary antibodies diluted in blocking buffer (1:100 anti-PYK2: ab32571, Abcam; 1:100 anti-Pax: AHO0492, ThermoFisher; 1:80 anti-phospho-PYK2 pY402: sc-293142, Santa Cruz; 1:80 anti-EEA1: GT10811, Invitrogen). The next day, cells were washed three times with PBS and incubated for 1 hour at room temperature in the dark with secondary antibodies (1:200). Cells were washed three times, and drops of ProLong^TM^ Glass Antifade Mountant with NucBlue^TM^ Stain (ThermoFisher, P36981) were applied to stain the nuclei and preserve the samples (long-term stored at -20°C). For plasma membrane staining Vybrant™ DiD Cell-Labeling Solution (V22887, Invitrogen) was added before imaging according to the manufacturer’s instructions. Images were acquired with Leica Stellaris 8 FALCON confocal microscope equipped with a Zeiss 63x/1.4 oil objective and processed on Leica LAS-X. Microscopy figures were edited with ImageJ (version 1.54c), maintaining the original pixel intensity range. For colorblind-friendly figures, images were converted to 16-bit, and lookup tables for green, magenta, and blue were selected.

### Fluorescence Recovery After Photobleaching (FRAP)

HEK293T cells transfected with different EGFP-PYK2 constructs were plated in glass-bottom dishes and imaged on an inverted Zeiss LSM 880 confocal microscope. EGFP was excited with a 488 nm laser, and the microscope was controlled with Zeiss Zen software. Images were acquired with a 63x/1.4 oil immersion objective with a 3x optical zoom. The droplets in cells were bleached for 10 seconds with a maximum laser power of a 488-nm laser (1 AU). The recovery was recorded for 60 time points after bleaching (60 seconds). The fluorescent intensity of the bleached area over time was calculated by Zen.

### Total Internal Reflection Fluorescence (TIRF) and epifluorescence microscopy

The day after transfection, HeLa cells were seeded onto 35 mm glass-bottom dishes pre-coated with low (20 µg/mL) or high (50 µg/mL) fibronectin^[16]^. After 6 hours, cells were washed with warm PBS and maintained in phenol red-free DMEM supplemented with 10% FBS for imaging. TIRF and epifluorescence imaging were performed using a Zeiss Elyra 7 microscope equipped with a Plan-Apochromat 63x/1.46 Oil Korr M27 objective and a 488 nm laser line for EGFP excitation. For TIRF, the evanescent field penetration depth was set to ∼100 nm to selectively excite fluorophores at or near the basal plasma membrane. Imaging was conducted at 37 °C in a humidified chamber with 5% CO₂ using a stage-top incubation system. Time-lapse sequences were acquired at 5-second intervals. Image acquisition and initial processing were performed using Zeiss ZEN Black software. Post-acquisition processing was carried out using Fiji (version 1.8.0_322), applying the “*rainbow RGB*” look-up table.

### Image analysis

Intensity profile plots were generated using the *Plot Profile* tool in ImageJ (version 2.14.0/1.54f). For each image, corresponding regions of interest were selected with the freehand line tool and analyzed as single-channel data (green or magenta). Scale was globally set according to the distance reported on the original acquisition. Quantitative analysis of the IACs was performed on ImageJ using the protocol reported by Horzum et al, 2014^[74]^. In brief, background was subtracted from the images and enhanced local contrast was run. Subsequently, the automatic threshold was applied and the particles analysed. For each sample, at least five independent acquisitions were evaluated. Only EGFP-positive cells were selected for the analysis of transfected cells.

### IDR database construction

We extracted disordered regions from the AlphaFold human database (v4) based on a restricted high-pLDDT fraction and a minimum length threshold. Specifically, sequences were divided into smaller fragments using 18 combinations of high-pLDDT percentage (≤10%, 15%, and 30% of residues with pLDDT > 80) and sequence length (100, 110, 120, 130, 140, and 1022 residue-long windows, with a 20-residue overlap). The resulting human IDR database comprised a total of 8,557,526 unique sequences. These sequences were subsequently used to calculate ESM-2 embeddings and assess their similarity to the PYK2 KFL sequence.

### Similarity scoring and IDR ranking

We calculated embeddings for the complete human IDR database using the esm2_t48_15B_UR50D model. The cosine similarity of each sequence embedding was then computed against the embedding of the PYK2 KFL (residues 728-839). Sequences were ranked based on their similarity scores.

### Bioinformatic analysis of IDR sequences

We computed various biophysical properties for the IDR sequence database using the CIDER and Biopython Python libraries. These properties are illustrated in **Suppl. Fig. 4**.

### GO enrichment analysis

From each of the 18 IDR groups described in the IDR Database Construction section (excluding the 1022 residue-long groups), we selected the 500 sequence fragments with the highest similarity to PYK2-KFL. The genes associated with these fragments were identified, resulting in a list of 150 common genes.

Gene Ontology (GO) enrichment analysis was performed using the DAVID tool^[75]^. This analysis yielded a list of enriched GO terms within the set of 150 genes. To summarise and visualise the GO terms and their associated P-values, we processed the results with REVIGO ^[76]^. The summarised results are presented in **Suppl. Fig. 5**.

### Statistical analysis

Graphs represent data from at least three independent experiments. For normally distributed datasets, results are presented as mean ± standard deviation (SD). Comparisons between two groups were performed using a two-tailed Student’s *t*-test. A *p*-value < 0.05 was considered statistically significant. All statistical analyses and graph plotting were performed using OriginPro (OriginLab, version 9.7).

## ACKNOWLEDGEMENTS

We would like to thank the Imaging and Characterization Core Lab at KAUST for the NMR, and confocal microscopy time. We also thank the Bioscience Core Lab at KAUST for the help with mass-spectrometry data collection and analysis. For computer time, this research used the resources of the KAUST Supercomputing Laboratory (KSL). We thank N. Kathiresan and G. Wickham from the KSL for their support. This publication is based on the work supported by the King Abdullah University of Science and Technology (KAUST) through the baseline fund and the Award No. URF/1/2602-01-01 from the Office of Sponsored Research (OSR), and the KAUST Center of Excellence for Smart Health (KCSH), under award number 5932. RH and MK acknowledge funding from the Near Term Grand Challenge Grant (REI/1/5235-01-01) and CRG 11 (URF/1/5041-01-01) from KAUST.

## AUTHOR CONTRIBUTIONS

Cell biology: I.S, G.C, P.Y, S.AF; Production of expression plasmids: G.K, J-A.G; Protein cloning, expression and purification: A.A.M, A.A; Confocal and super-resolution microscopy: G.C; Biochemical and biophysical analysis: A.A.M, A.A, S.T.A; NMR analysis: A.A.M, K.S, S.AH, Ł.J, S.T.A; Computational analysis: F.J.G.V, M.K, R.H; Project design and supervision: S.T.A, A.A.M, I.S, Ł.J; Manuscript writing (draft, coordination): S.T.A. All authors have contributed text for the manuscript, commented and approved the manuscript.

