## Supplementary material for "Molecular Basis and Cellular Effects of Janus-Class–Driven Cytoplasmic PYK2 Coacervates": Suppl. Material

| Condition | Condition | Condition |
|---|---|---|
| 300 µM; 200mM NaCl; 25°C | 100 µM; 200mM NaCl; 25°C | **50 µM; 200mM NaCl; 25°C** |
| 300 µM; 500mM NaCl; 25°C | 100 µM; 500mM NaCl; 25°C | 50 µM; 500mM NaCl; 25°C |
| 300 µM; 1000mM NaCl; 25°C | 100 µM; 1000mM NaCl; 25°C | 50 µM; 1000mM NaCl; 25°C |

**Suppl. Table. 1**
LLPS droplets visualised by fluorescence microscopy of Alexa 488-labelled $KFL_{728\text{-}839}$ under variable protein and NaCl concentrations. Red – No LLPS.

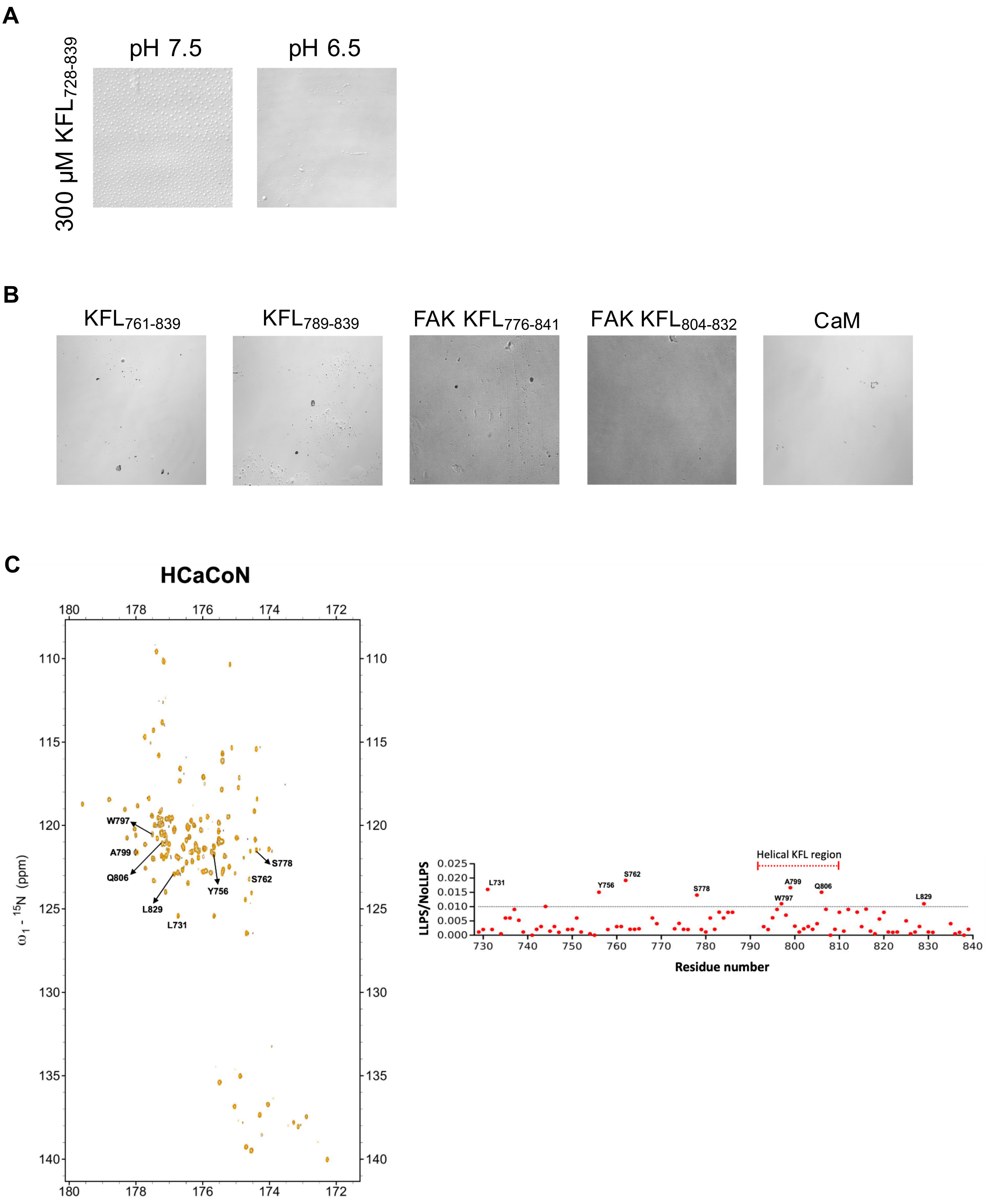


**Suppl. Fig. 1:**

**A.** Non-labeled $KFL_{728\text{-}839}$ visualized by DIC showing droplet formation at pH 7.5, and no droplet at pH 6.5.

**B.** Non-labeled $KFL_{761\text{-}839}$, $KFL_{789\text{-}839}$, FAK-$KFL_{776\text{-}841}$, FAK-$KFL_{804\text{-}832}$, and CaM visualized by DIC showing no droplet formation.

**C.** The LLPS formation process does not affect the backbone conformation and dynamics of PYK2. Left panel is the $^{13}C$ detected CON spectrum of PYK2 under LLPS conditions and non-LLPS conditions. Gold represents LLPS of PYK2 while black represents non-LLPS conditions for PYK2. Residues exhibiting $^{13}C/^{15}N$ CSP higher than noise (median + 1.5 IRQ) are highlighted.

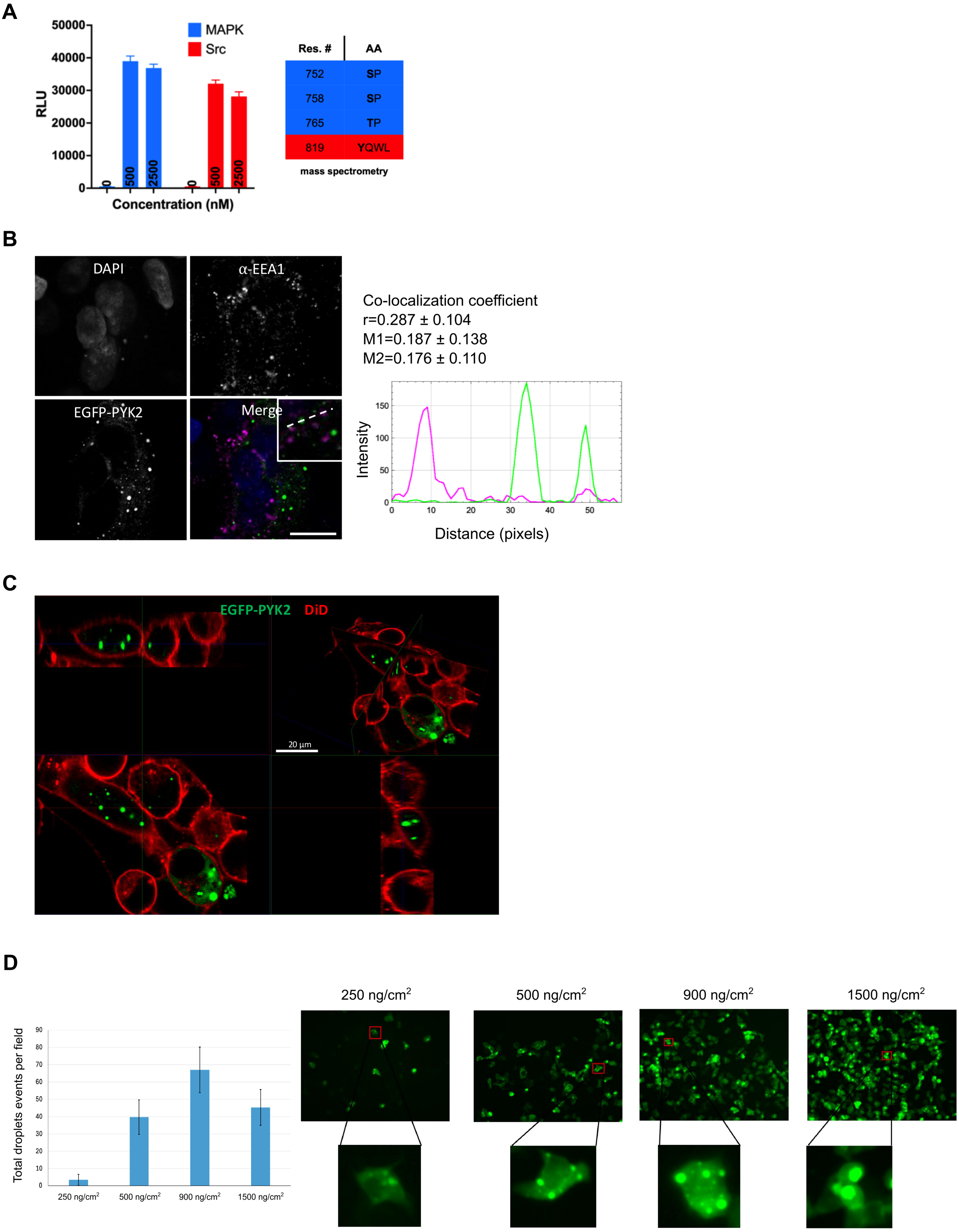


**Suppl. Fig. 2:**

**A.** Correlation of % ATP conversion to ADP using the ADP-Glo™ assay for $KFL_{728\text{-}839}$ in the absence (0 nM) and presence of (500 nM) and (2500 nM) of MAPK (blue) and Src (red). (Right) Table showing phosphorylated residues in the presence of MAPK (blue rows) and Src (red rows) identified using mass spectrometry.

**B.** (Left) Single-channel and merged immunofluorescence images of HeLa cells transfected with EGFP-PYK2 (green), stained anti-EEA1 antibody (magenta) and counterstained with DAPI (blue). (Right) Co-localization analysis including profile plots and Pearson/Manders coefficients calculated from n = 5. Scale bar: 10 μm. Images were acquired with Leica Stellaris 8 FALCON confocal microscope equipped with a Zeiss 63x/1.4 oil objective.
**C.** Side views of confocal images of HeLa cells transfected with EGFP-PYK2 (green) and stained with DiD (red). Scale bar: 20 μm. Images were acquired with Leica Stellaris 8 FALCON confocal microscope equipped with a Zeiss 63x/1.4 oil objective.
**D.** (Left) Bar plot showing the number of PYK2 droplets per cell varying the amount of DNA transfected. (Right) Representative fluorescence images for each condition. Images were acquired with FLoid™ Cell Imaging Station.

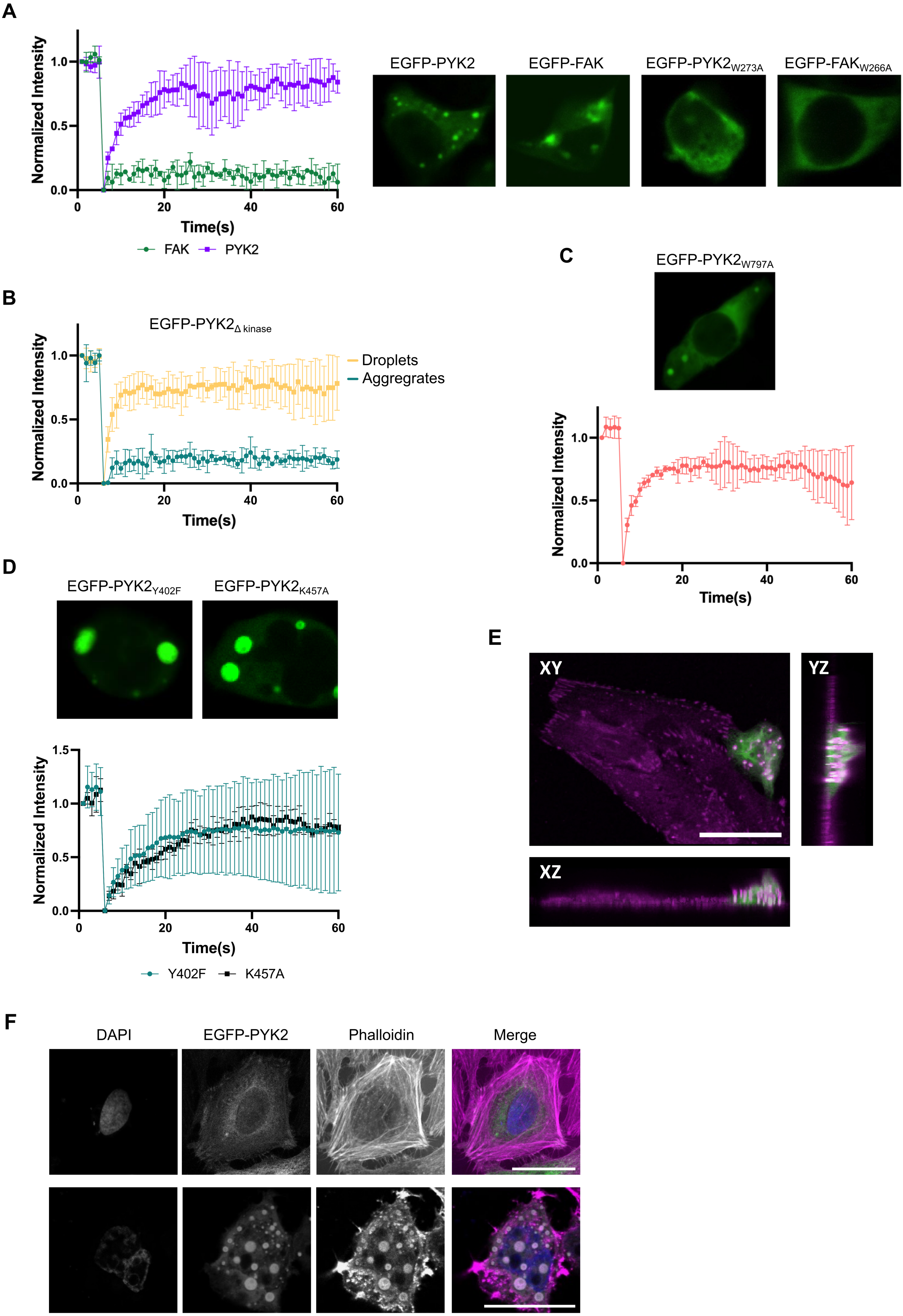

A
Normalized Intensity
Time(s)
FAK
PYK2
EGFP-PYK2
EGFP-FAK
EGFP-PYK2W273A
EGFP-FAKW266A
B
EGFP-PYK2Δ kinase
Droplets
Aggregrates
C
EGFP-PYK2W797A
D
EGFP-PYK2Y402F
EGFP-PYK2K457A
Y402F
K457A
E
XY
YZ
XZ
F
DAPI
EGFP-PYK2
Phalloidin
Merge

**Suppl. Fig. 3:**
**A.** (Left) FRAP curves showing the recovery of EGFP-FAK and EGFP-PYK2 puncta after photobleaching. Data are presented as mean and standard error (n=3). (Right) Representative images of HEK293T cells transfected with EGFP-PYK2, EGFP-FAK, EGFP-PYK2$_{W273A}$ and EGFP-FAK$_{W266A}$. Images were acquired with FLoid™ Cell Imaging Station.
**B.** FRAP curves of EGFP-PYK2$_{\Delta kinase}$ forming either droplets that recover (yellow) or static aggregates (green). Data are presented as mean and standard error (n=3).
**C.** (Top) HEK293T transfected with EGFP-PYK2$_{W797A}$ and (bottom) FRAP curve. Data are presented as mean and standard error (n=3). Images were acquired with FLoid™ Cell Imaging Station.
**D.** (Top) HEK293T transfected with EGFP-PYK2$_{Y402F}$ and EGFP-PYK2$_{K457A}$. (Bottom) FRAP curves. Data are presented as mean and standard error (n=3). Images were acquired with FLoid™ Cell Imaging Station.
**E.** Side views of confocal images of HeLa cells transfected with EGFP-PYK2 (green) and stained with anti-paxillin antibody (magenta). Scale bar: 20 µm. Images were acquired with Leica Stellaris 8 FALCON confocal microscope equipped with a Zeiss 63x/1.4 oil objective.
**F.** Low (top row) and high (bottom row) EGFP-PYK2 (green) expression in HeLa cells. Actin stained with phalloidin (magenta) and counterstaining with DAPI (blue). Scale bar: 10 µm. Images were acquired with Leica Stellaris 8 FALCON confocal microscope equipped with a Zeiss 63x/1.4 oil objective.

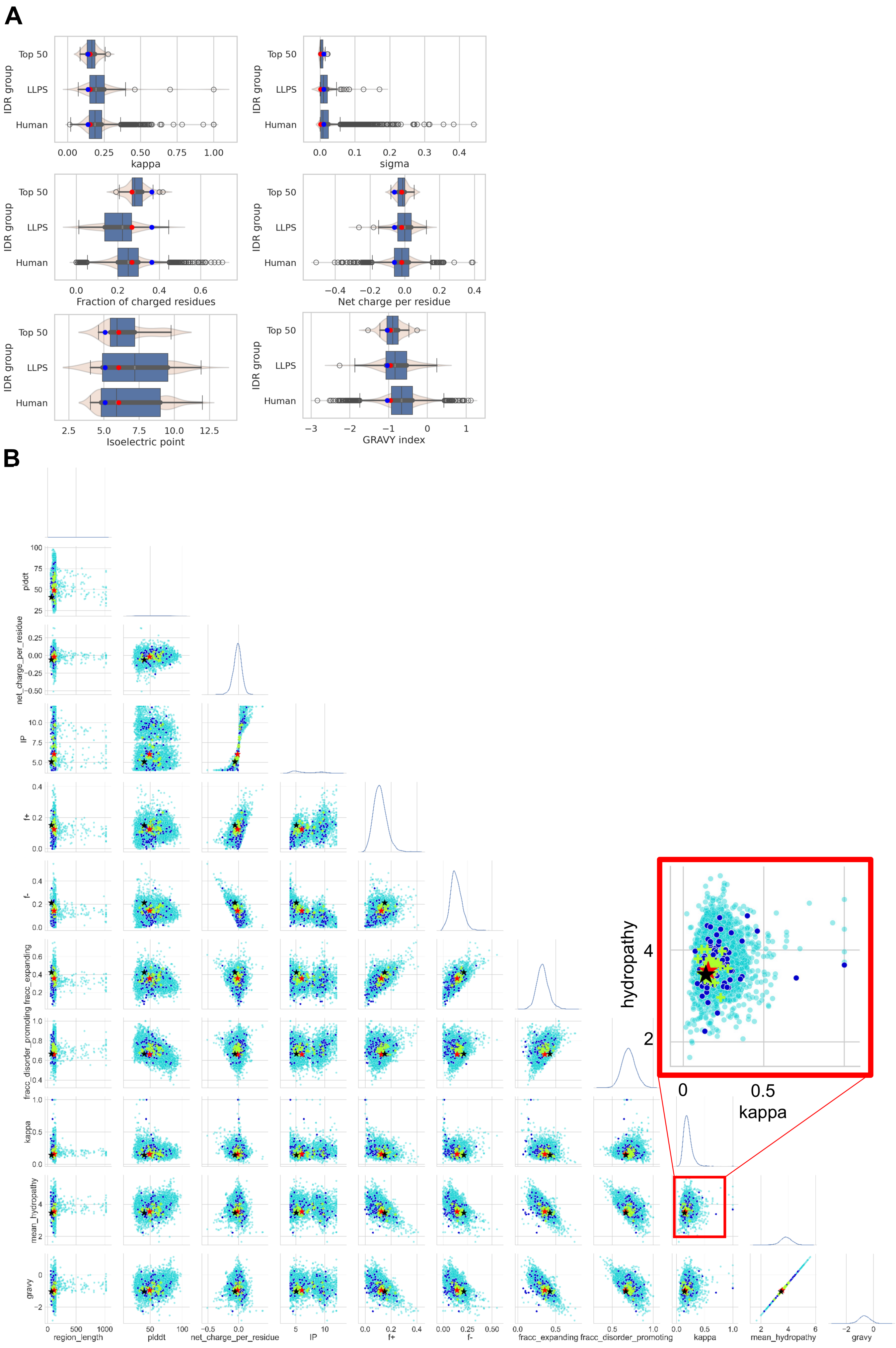
A
IDR group
Top 50
LLPS
Human
kappa
sigma
Fraction of charged residues
Net charge per residue
Isoelectric point
GRAVY index
B
plddt
net_charge_per_residue
IP
f+
f-
fracc_expanding
fracc_disorder_promoting
kappa
mean_hydropathy
gravy
region_length
hydropathy
kappa

**Suppl. Fig. 4:**
**A.** Biophysical properties for the three groups of IDRs. In every case, the group with the most similar sequence embeddings to PYK2 KFL (Top 30) is closer in the values of the properties. The LLPS driver sequences and the human IDRs have a higher variation.
**B.** This pattern is consistent when plotting the 10 studied properties on the 2D plots on panel.

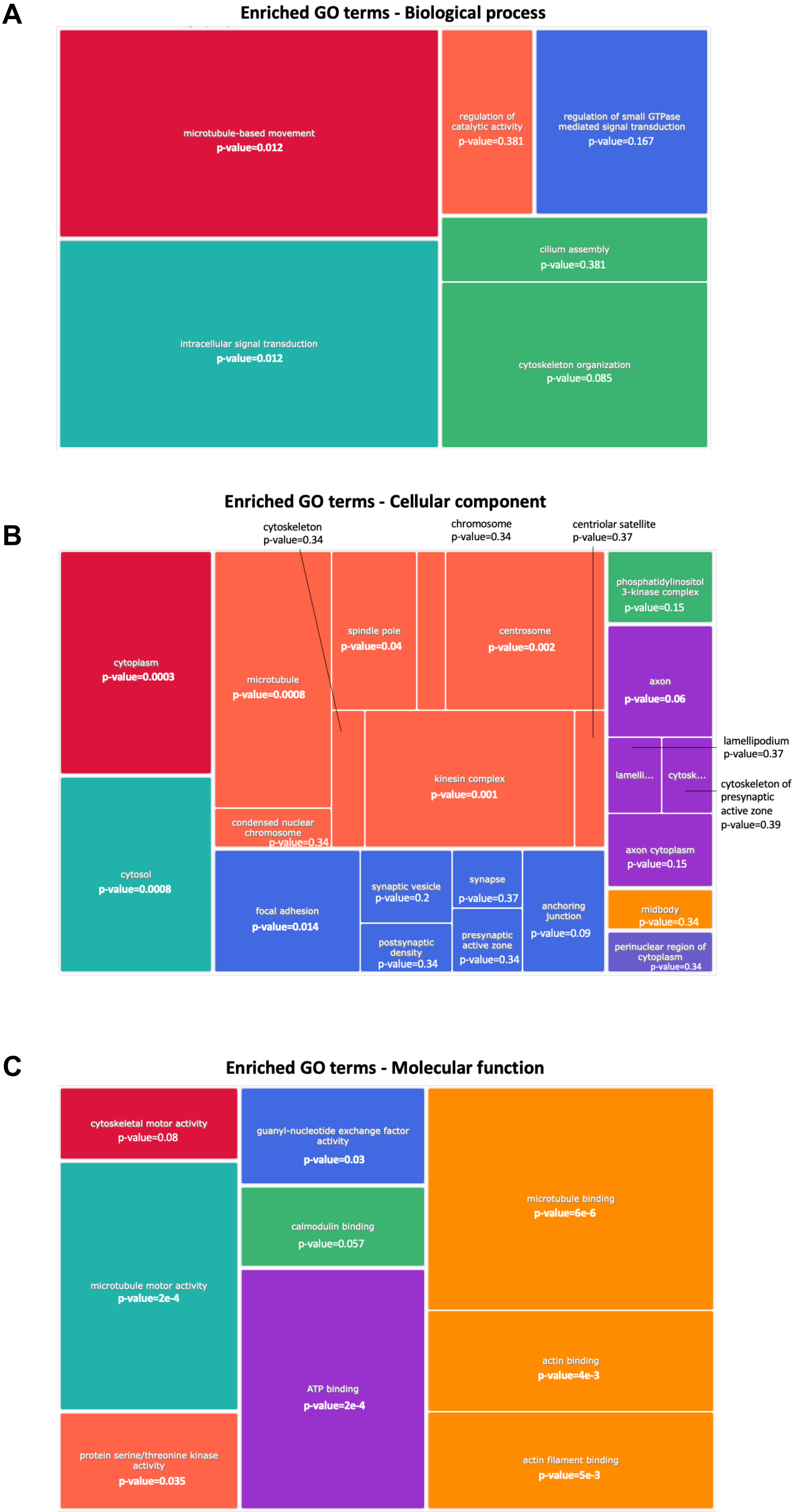

A
Enriched GO terms - Biological process
microtubule-based movement
p-value=0.012
regulation of catalytic activity
p-value=0.381
regulation of small GTPase mediated signal transduction
p-value=0.167
cilium assembly
p-value=0.381
intracellular signal transduction
p-value=0.012
cytoskeleton organization
p-value=0.085
B
Enriched GO terms - Cellular component
cytoskeleton
p-value=0.34
chromosome
p-value=0.34
centriolar satellite
p-value=0.37
cytoplasm
p-value=0.0003
microtubule
p-value=0.0008
spindle pole
p-value=0.04
centrosome
p-value=0.002
phosphatidylinositol 3-kinase complex
p-value=0.15
axon
p-value=0.06
lamellipodium
p-value=0.37
lamelli...
cytosk...
cytoskeleton of presynaptic active zone
p-value=0.39
kinesin complex
p-value=0.001
condensed nuclear chromosome
p-value=0.34
cytosol
p-value=0.0008
focal adhesion
p-value=0.014
synaptic vesicle
p-value=0.2
synapse
p-value=0.37
anchoring junction
p-value=0.09
postsynaptic density
p-value=0.34
presynaptic active zone
p-value=0.34
axon cytoplasm
p-value=0.15
midbody
p-value=0.34
perinuclear region of cytoplasm
p-value=0.34
C
Enriched GO terms - Molecular function
cytoskeletal motor activity
p-value=0.08
guanyl-nucleotide exchange factor activity
p-value=0.03
microtubule binding
p-value=6e-6
calmodulin binding
p-value=0.057
microtubule motor activity
p-value=2e-4
actin binding
p-value=4e-3
ATP binding
p-value=2e-4
protein serine/threonine kinase activity
p-value=0.035
actin filament binding
p-value=5e-3

**Suppl. Fig. 5:**
**A.** Summarization of enriched GO terms from 150 genes with highly similar sequence embeddings to PYK2-KFL.

**Suppl. Video 1:** EGFP- tagged PYK2 droplets fuse over time in cells. HeLa cells were transfected with EGFP-PYK2 plasmid and live-cell imaging was recorded for 45 seconds with a Leica Stellaris 8 FALCON confocal microscope with a Zeiss 63x/1.4 oil objective and processed using Leica LAS-X software.

**Suppl. Video 2:** EGFP-PYK2 droplets visualized by TIRF microscopy.HeLa cells were transfected with an EGFP-PYK2 plasmid, and live-cell imaging was performed using total internal reflection fluorescence (TIRF) microscopy on a Zeiss Elyra 7 microscope equipped with a Plan-Apochromat 63×/1.46 Oil Korr M27 objective.

**Suppl. Video 3:** EGFP-PYK2 droplets visualized by epifluorescence. HeLa cells were transfected with an EGFP-PYK2 plasmid, and live-cell imaging was performed using epifluorescence with Zeiss Elyra 7 microscope equipped with a Plan-Apochromat 63x/1.46 Oil Korr M27 objective.
